# IMPRESS: AI-coupled HPC workflows for protein design

**DOI:** 10.64898/2026.11.13.766417

**Authors:** Mason Hooten, Amanda Liyanaarachchi, Aymen Alsaadi, Mariya Goliyad, Shantenu Jha, Sagar D. Khare

## Abstract

Protein design campaigns are often run as multi-stage pipelines with a generative backbone model, an inverse-folding model, and a structure predictor chained together in that order. The stages differ by one to two orders of magnitude in cost per candidate, and the allocation of sampling effort across them is fixed before a campaign begins. Such a filter cascade can only discard candidates. It cannot redirect sampling toward the regions of design space a model stack can reach, and for targets whose constraints are far from the generative model’s training distribution it may return no solution at any practical sample count. We present IMPRESS (Integrated Machine-learning for PRotein Structures at Scale), a framework in which design pipelines are stateful objects that halt, continue, resample, or branch at runtime from their own accumulated results. Adaptive decisions are evaluated asynchronously with respect to execution, so a campaign refines itself without global synchronization. We apply IMPRESS to three problems: backbone generation on discontinuous enzyme active-site motifs, *de novo* small-molecule binder design, and PDZ binder redesign. On an enzyme active-site benchmark, adaptive parameterization solves 17 of the 29 motifs carried on three or more discontinuous residue islands against a nonadaptive reference’s 8, with no motif solved by the reference alone. In the small-molecule case the adaptive rules shift the realized task mix toward inexpensive sequence exploration and away from expensive folding. In the interface redesign case the adaptive pipelines sample more broadly at each sequence position while keeping each input’s ensemble closer together, so exploration is targeted rather than random. All comparisons reported here are between *in silico* metrics. We expect that adaptive parameterization will matter most for targets furthest from the training distributions of the underlying models, and that the same primitives will accommodate experimental measurement as a steering signal.

**TOC Graphic:** 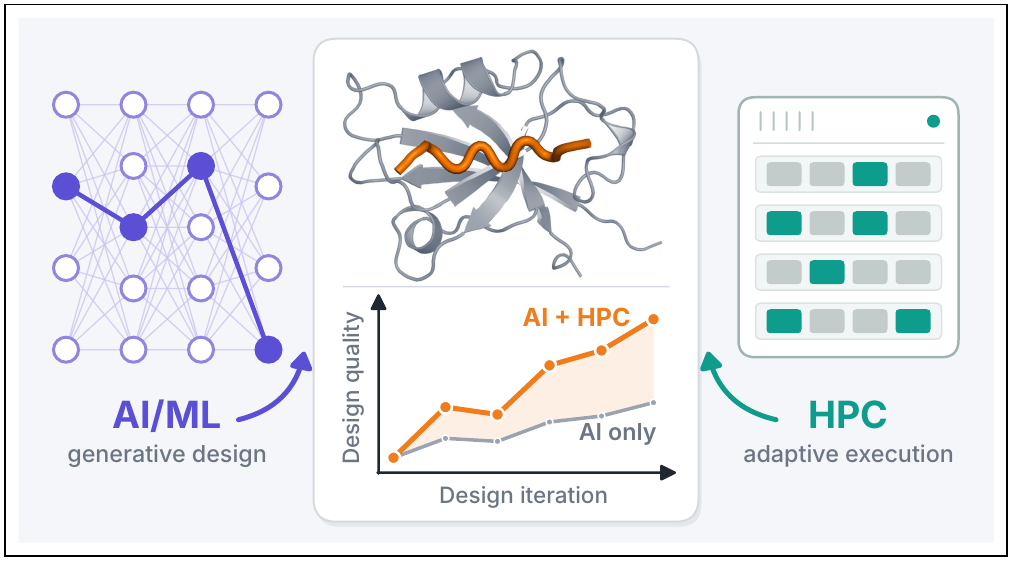

## 1 Introduction

Protein design workflows are often implemented as a pipeline. A diffusion model generates a backbone, an inverse-folding model designs a sequence onto that backbone, and a structure predictor folds the sequence to check that it returns the intended geometry.^1–3^ The same three-stage approach serves targets as different as a protein interface, a small molecule, and an enzyme active site,^4–6^ and each step is individually sufficiently reliable, and computationally efficient, that campaigns of 10^3^ to 10^5^ candidates are routine. Thus the binding constraint on a design campaign is not the accuracy of any single model, but rather how sampling effort is distributed across the stages.

The distribution of sampling effort is almost always fixed before the campaign starts. A typical protocol generates a set number of backbones (e.g. 1000–10000), designs a set number of sequences per backbone (5–10), folds each sequence, and retains whatever survives a set of confidence and geometry filters.^4,7^ Thus, the campaign is a filter cascade with constant fan-out. This is a sensible default, and the filtering is essential because the stages differ by roughly one to two orders of magnitude in cost per candidate: inverse folding is seconds, backbone diffusion is tens of seconds, and all-atom structure prediction is minutes-hours.^2,8,9^ Discarding a candidate before it reaches the structure predictor is therefore the single largest economy available to a fixed pipeline.

However, a filter cascade can only remove candidates, it cannot propose better-placed ones. Where the design constraint falls within or close to what the generative model produces readily, this costs little, and raising the sample count leads to a roughly proportional number of hits. However, in cases where the constraint sits far from the model’s training distribution, it costs a great deal. Enzyme active sites are the classic case: the catalytic residues are fixed in identity and in relative geometry, and they are typically carried on three or more discontinuous segments of sequence, so the generative model is asked to satisfy several simultaneous point constraints that it rarely satisfies by chance.^6,10–12^ Reported success rates on such targets are low, and for the hardest of them they are zero at any sample count that is practical to run.^12,13^ Adding samples to a cascade that filters all of the proposed solutions out is therefore highly ineffective.

Pipeline filters are usually the models’ own confidence estimates, and those estimates are calibrated on natural structures rather than on designs.^14^ Idealized de novo backbones can score very highly and still fail experimentally, while unusual but correct geometries can score poorly.^7,15^ Chaining several such models compounds the effect, since each stage inherits the preceding stage’s bias and adds its own. Thus, for pipeline construction, a filtering threshold fixed before any results exist is a poor instrument for locating the regions of design space a given stack can actually reach, as such a threshold lacks context of the campaign it is embedded in.

To resolve high-quality seams in the joint inference landscape, pipeline parameters must be set in the context of joint measurements of a reference ensemble. Methods that contextualize a design in relation to the ensemble, such as statistical ensemble comparisons or surrogate models, may be used to steer pipeline task parameterization. Adaptive steering can be used to strategically explore the inference space, resolving high-quality regions of the design space over tunable pipeline parameters.

Reallocating sampling effort at runtime is a well-developed idea in molecular simulation. Adaptive seeding, FAST, and REAP each choose where to spend the next batch of simulation using the ensemble accumulated so far,^16–18^ and DeepDriveMD and Colmena generalize this to learned surrogates steering ensembles at scale.^19–21^ Those systems steer one sampler against one progress coordinate. A design campaign is a harder target. The stages are different models with different failure modes, different costs, and scores that are not commensurable with one another, so what must be carried forward is not a single coordinate but the pipeline’s own accumulated record of backbones, sequences, and folds.

Here we present IMPRESS (Integrated Machine-learning for PRotein Structures at Scale), a framework for expressing design campaigns whose task allocation is determined at runtime by their own intermediate results. An IMPRESS workflow is written as a set of stateful pipelines. Each pipeline holds its own history and reaches explicit decision points at which it may halt, continue, resample a previous stage, or spawn child pipelines that inherit the parent’s state and then adapt independently. Decisions are evaluated asynchronously with respect to execution, so a campaign refines itself without a global synchronization barrier and tasks with very different resource profiles are kept in flight together. This allows the exploration to expand both in the number of candidates considered and in the number of adaptive paths followed. The framework extends our earlier adaptive design middleware^22^ and executes through RADICAL-AsyncFlow^23^ and the RHAPSODY AI-HPC runtime,^24^ which place heterogeneous inference and simulation tasks on heterogeneous resources.^25^

We apply IMPRESS to three design problems that between them span the adaptive moves available to a pipeline. The first scaffolds backbones onto discontinuous enzyme active-site motifs, accepting a failed design piecewise by fixing the parts of it that passed and re-posing the remainder as a new problem. The second designs de novo small-molecule binders under adaptive quality gates that limit the work done on low-scoring design intermediates and a greedy strategy that forwards a high-confidence fold to the next iterative diffusion stage as a scaffold. The third redesigns a protein-peptide interface with a genetic algorithm that adapts in sequence space alone.^26,27^ We show that runtime reallocation changes the region of design space a campaign can reach, and separately reshapes how a campaign distributes work across the stages. On the active-site benchmark, restricted to the 36 motifs both campaigns folded, the adaptive workflow solves 17 of the 29 motifs carried on three or more residue islands against the nonadaptive reference’s 8, under a matched per-motif fold budget and with no motif solved by the reference alone (exact McNemar *p* = 0.002). We read this as evidence of reach rather than of economy, since the adaptive campaign is not a cheaper route to the same designs but a route to designs the reference does not produce. Because the reference campaign recorded every acceptance criterion without acting on any of them, the behaviour of a filtered but nonadaptive cascade over the same sample can be derived from the retained data. Doing so discards a fifth of the fold budget and solves no additional motif in any stratum, so gating accounts for none of the difference. In the small-molecule case, the adaptive rules shift the realized task mix toward inexpensive sequence exploration and away from expensive folding. For the interface redesign problem, the adaptive pipelines sample a broader set of residues at each sequence position while keeping each input’s ensemble closer together, so exploration is targeted rather than random, aiding the curation of a starting library for experimental evaluation.

The IMPRESS software is publicly available on Github^28^ https://github.com/radical-collaborati IMPRESS. The repository contains the code and data for the experiments described in this work along with documentation and interactive tutorials.

### 1.1 Tasks in generative protein design

The three use cases share computational motifs and a common adaptive vocabulary. The calculations typically use generative ML models to infer backbones *de novo*, sequences from backbones, and folded structures from sequences.^1,2,8^ The different inference modes are effectively independent, and thus each admits its own techniques for constraining the joint design space. In adaptive workflows, the parameterization of a task depends on the results of preceding tasks. In ML models, reparameterization may mean additional training (e.g. finetuning on a domain-specific sample or reinforcing on a growing set of ‘hit’ designs), or refining the invocation of the model (i.e. the parameterization of its API) to steer sampling in response to the accumulating ensemble of design candidates.

Although many workflow designs are based on inference in the space of physical features, latent-space simulation pipelines are also possible. Thus, a workflow may be designed to chain inference at different scales or in transformed representations, and this flexibility maximizes control over the design space. Multiscale campaigns may target combinations of physical and black-box scores, targeting e.g. improved scores or reduced variance on the desired metrics. The use cases below demonstrate these capabilities along with the attendant reshaping of the computational load that adaptive sampling can impart.

### 1.2 Related Work

Recent surveys map the available tooling onto the steps of a design workflow and guide their assembly end to end;^29^ what they organize is composition. Workflow systems have largely settled on the directed acyclic graph as the unit of that composition, from general engines such as Nextflow^30^ to protein-specific frameworks. BioPipelines is the latter case, placing over forty generation, design, prediction, and analysis tools behind standardized biomolecular streams, so that a protocol written as a short Python script prototypes in a notebook and submits to a cluster unchanged.^31^ Its separation of configuration from execution is the architectural contrast with IMPRESS. Configuration runs in Python, predicts the file-system structure the run will produce, and emits self-contained shell scripts; execution follows only after the Python process has terminated, so no orchestrator persists during the run. Values computed at runtime do propagate between cycles—in their binding-pocket example the redesigned positions are recomputed each cycle from the current best structure—but the task graph is materialized in full before anything executes, and a loop is unrolled to a cycle count fixed in the script. That trade buys inspectable, independently resubmittable artifacts. IMPRESS trades the other way: a resident manager holds per-pipeline state, so the graph is a runtime product rather than an artifact of submission, and a pipeline may halt, resample, or branch on evidence that did not exist at submission.

A second class packages one protocol end to end. BindCraft hallucinates binder sequences by backpropagating a composite loss through AlphaFold2 weights, optimizes the non-interface core with ProteinMPNN, repredicts and relaxes, then screens against nine thresholds on model confidence, interface shape complementarity, hydrogen bonding, and surface hydrophobicity.^32^ Those thresholds were set from experimental observations in earlier campaigns and refined over the course of the work, and the pipeline reports experimental success rates of 10–100%. The thresholds are constants, fan-out is constant at twenty sequences per trajectory with at most two admitted, and the campaign loops until a target count of passing designs accumulates. Such a cascade discards candidates but cannot propose better-placed ones, and when the constraints sit far from the generative models’ training distribution the filters may admit nothing at any practical sample count. IMPRESS is not a competing protocol but a way of expressing protocols whose thresholds and routing are set against the ensemble as it accumulates.

A third class delegates orchestration to language models. ChemGraph decomposes a natural-language request with a planner agent, steps the subtasks through executor agents in a reason-and-act loop, and aggregates the results, with graph neural network foundation models supplying the underlying calculations;^33^ ProtAgents runs planner, assistant, and critic agents in a dynamic group chat over physics-based and learned models.^34^ What adapts there is the plan—which tool to invoke, and how to recover when one errors—assessed against reference answers specifying the expected tool calls and result for a single request. In IMPRESS the transformations are fixed and their parameterization and routing adapt, on numerical ensemble-relative evidence, across a population rather than a request. The two compose rather than compete: an IMPRESS decision point may itself invoke an LLM-based reasoning component (Section 3).

Runtime reallocation of sampling effort is established in molecular simulation, where adaptive seeding, FAST, and REAP select the next batch from the accumulated ensemble,^16–18^ and DeepDriveMD and Colmena generalize this to learned surrogates steering ensembles at scale,^19–21^ now a recognized family of AI-coupled HPC motifs.^25^ These steer one sampler against one progress coordinate. A design campaign chains models of differing cost and failure mode whose scores are not commensurable, so what carries forward is the pipeline’s record of backbones, sequences, and folds rather than a scalar coordinate. IMPRESS makes that record a first-class per-pipeline object and evaluates routing against it asynchronously, without a global synchronization barrier.

Our work is distinguished from task-placement layers such as Parsl and Balsam, which likewise execute dynamic task graphs on HPC resources and which an IMPRESS-like system could in principle be built on.^35,36^ Those layers decide *where and when* a task runs, and the application decides *which* tasks exist; what IMPRESS and the RHAPSODY runtime beneath it add is that the task composition of the workflows is decided made at runtime, from the campaign’s own accumulated results, and that the two are decoupled – a pipeline’s adaptive function is evaluated while other pipelines continue executing, so refining the task set costs no barrier. The runtime contribution is narrow: heterogeneous inference and simulation tasks, with very different device and residency profiles, are kept in flight together under one placement policy, which is what makes a campaign of thousands of short, unequal tasks feasible without the application managing device allocation itself.^24,25^ An executor that places a fixed graph well does not supply either property.

The prior work thus separates on two axes: what a system composes, and what it permits to change once the campaign is running. Composition layers and packaged protocols are strong on the first and fixed on the second; agentic planners vary the plan but adapt on linguistic evidence; adaptive sampling frameworks adapt on numerical evidence but assume a single sampler and a single coordinate. IMPRESS occupies the remaining position, applying ensemble-relative adaptation to a heterogeneous, multi-stage design pipeline.

## 2 Use Cases and Motivating Workflows

We design three adaptive workflows targeting model use cases in protein design, composing tasks typical of ML-based protein design workflows.^37^ Many other workflow compositions are possible, with a broad and growing range of protein design models^37^ and leveraging a variety of compositional motifs.^25,29^

The first workflow addresses diffusion over discontinuous scaffolds, employing iterative definition of the design target to achieve solutions piecewise. The second workflow produces small-molecule binders using an explore/exploit strategy. Pipelines are initialized with back-bones generated *de novo* to target atoms on the ligand. Backbone-sequence designs yielding high-confidence folded structures are flagged as an external signal of sampling quality in the backbone generative model. The high-confidence folds are fed forward as a scaffold for a next round of generative diffusion, while low-confidence folds are not iterated and the workflow reverts to a *de novo* backbone. The third workflow redesigns a peptide binder given as a wild-type experimental structure. A genetic algorithm refines the target sequence on the given backbone, adapting only in the sequence space.

Each of the three workflows implements iterative design with conditional branching and adaptive parameterization. In addition to the design elements highlighted above, all of the workflows also employ judicious score gating at ML inference steps to mitigate wasteful computation, and as much as possible to keep the pipeline full of high-confidence candidates.

### 2.1 Backbone generation on discontinuous scaffolds

Enzyme design can be simplified in cases where the catalytic residues from a reference enzyme are already known from an experimental structure.^10,11^ Given a structure of the known enzyme-ligand binding pose, the atoms of the catalytic residues can scaffold a redesign workflow. In naturally occurring proteins, catalytic binding motifs typically involve residues far from each other in sequence space, where spatial coordination of the motif is mediated by conformation.^38^

Selecting a minimal subset of the binding motif as the design scaffold makes it possible to design a protein that is structurally distinct from the reference crystal structure. How-ever, scaffolding backbone generation with more than one or two discontinuous regions of the sequence, termed residue islands, can cause problems for diffusion models.^6,12^ Decoys generated on such discontinuous scaffolds using diffusion-based pipelines tend to fail common structural heuristic checks like motif RMSD leading to very low pipeline hit rates.^6,13^ Such failures may be attributed to a mismatch between the (inalterable) design constraints and the training of the model. To resolve this problem, it may be possible to iteratively compute a design with piecewise acceptance criteria, where good parts of a *de novo* candidate design are incorporated as fixed elements in successive iterations.

The discontinuous scaffolds workflow (Figure 2) begins with the generation of a backbone scaffolded on a given binding motif, with atoms of both the binder and target present. The workflow targets a pre-defined threshold for the quality of the predicted binding motif structure measured as RMSD. A structure with a low motif RMSD passes, and execution stops. When a structure fails, its positive features are used to refine the motif scaffold for a new pipeline.

**Figure 1:**
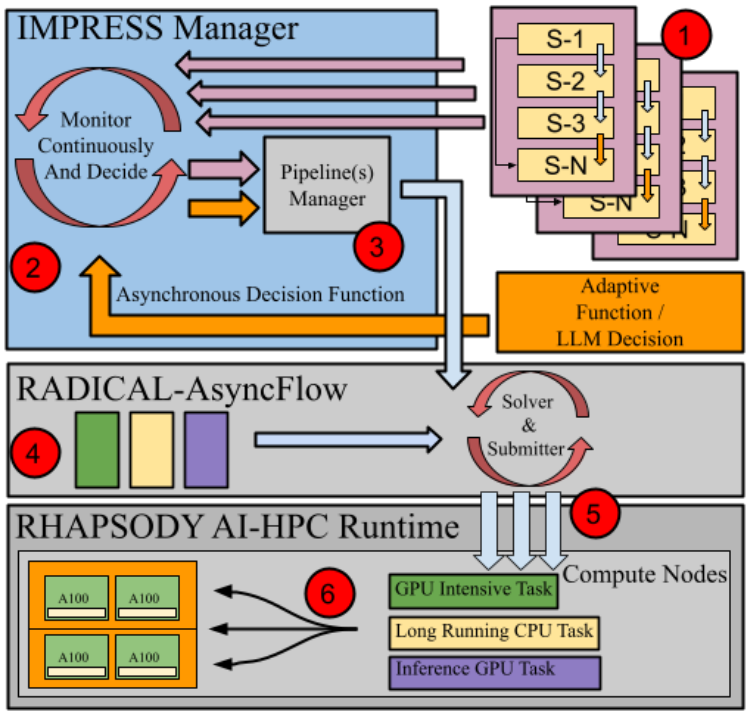
IMPRESS layered architecture and adaptive execution model. Task pipelines (1) are instantiated lists of tasks that eventually encounter adaptive logic gates. The results of adaptive logic are passed as a control signal to the IMPRESS Manager (2) which handles the ongoing task backlog and schedules tasks via the Pipeline Manager (3). AsyncFlow provides an asynchronous task runner loop (4), passing tasks to the runtime for execution (5), and listening for their resolution. The RHAPSODY runtime itself deploys tasks to the identified compute hardware (6) via pluggable backends. Choice of backend determines compatibility of the framework with a given compute resource, so that the availability of a compatible backend is the only framework-level dependency for IMPRESS.

**Figure 2:**
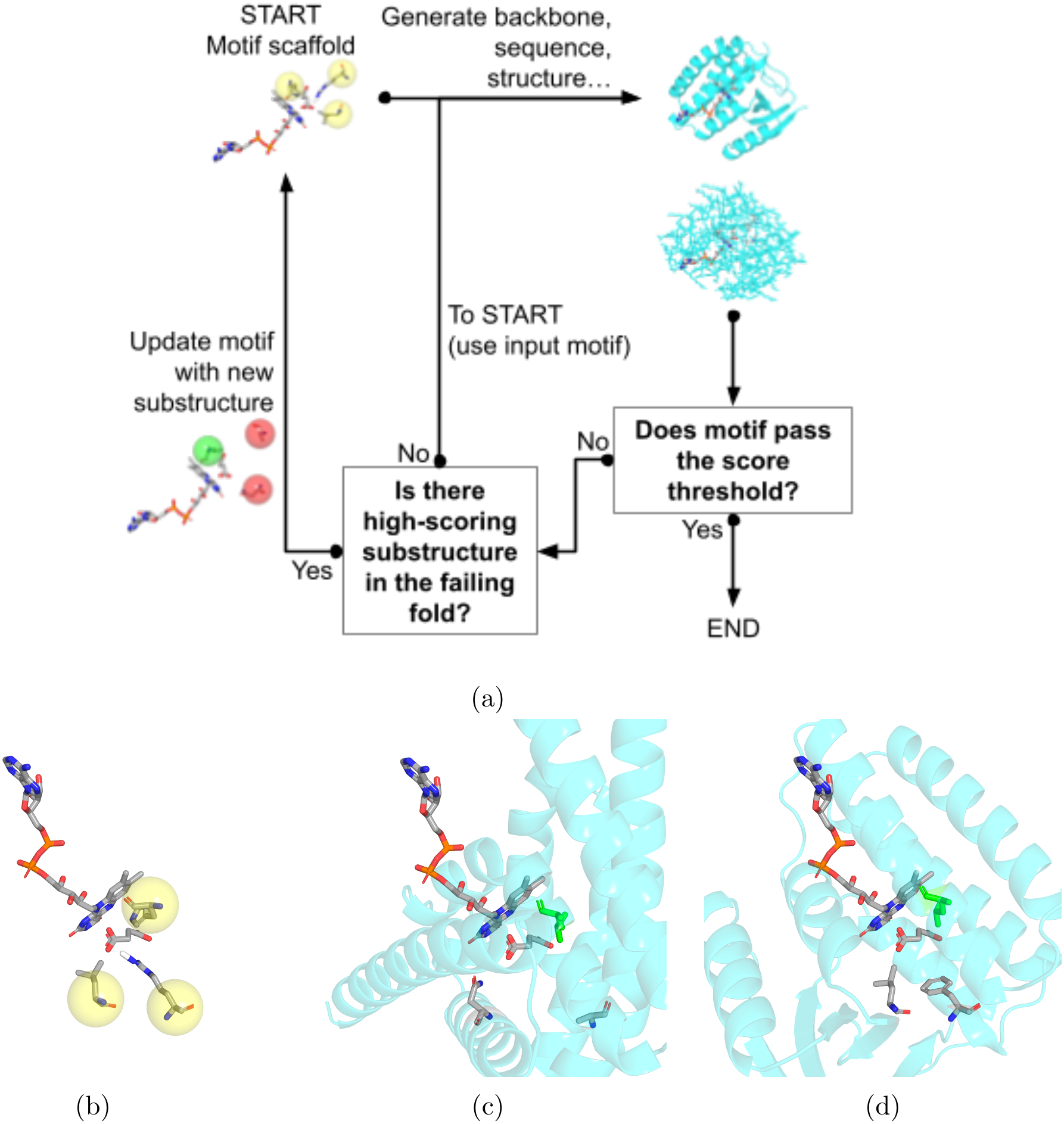
(a) The discontinuous scaffolds workflow. (b) A joint scaffolding motif comprising a small molecule and hot spot locations on the target protein. Protein hot spots indicated with yellow spheres. (c) The motif and surrounding context in the initial design. The green licorice element represents an anchor residue that survived iteration to contribute to the final design. (d) The motif and surrounding context in the final design. The anchor residue shown in green maintains its location and identity from the failed initial design, whereas the other two hot spots (the lower-most licorice segments in (c) and (d)) are revised by the intervening redesign steps.

The goal is to produce designs whose predicted structures closely recapitulate the atomic positions of the input binding motif. A backbone-sequence-structure pipeline is once again applied to translate the input binding motif scaffold into a predicted bound structure. Structures passing a tight threshold for RMSD of the binding motif are called a success, and the workflow stops. If a structure fails to pass the threshold, then motif atoms passing more stringent tests of local structure quality are identified and their coordinates are fixed, along with the identities and coordinates of intervening sequence segments. These fixed elements are used to reparameterize the input scaffold, allowing the workflow to solve the design piecemeal.

### 2.2 *De novo* small-molecule binders

Small-molecule ligands are an abundant target for binder applications in water treatment, energy, and biomedicine.^5,39^ Often the target molecule has no known binder, so designs are from scratch. A target site on the small molecules may be derived from its shape (as in the lock-and-key model of binding) or from relevant surface chemistry,^39,40^ and this is taken as the input constraint for *de novo* design.

The small-molecule binder workflow (Figure 3) produces designs targeting a small molecule, beginning with *de novo* generation of a suitable backbone, followed by sequence generation and fold prediction. Quality gates at each step combine fixed thresholds with comparisons against the growing ensemble of designs; Table 1 records which of them were active in the reported campaigns.

**Figure 3:**
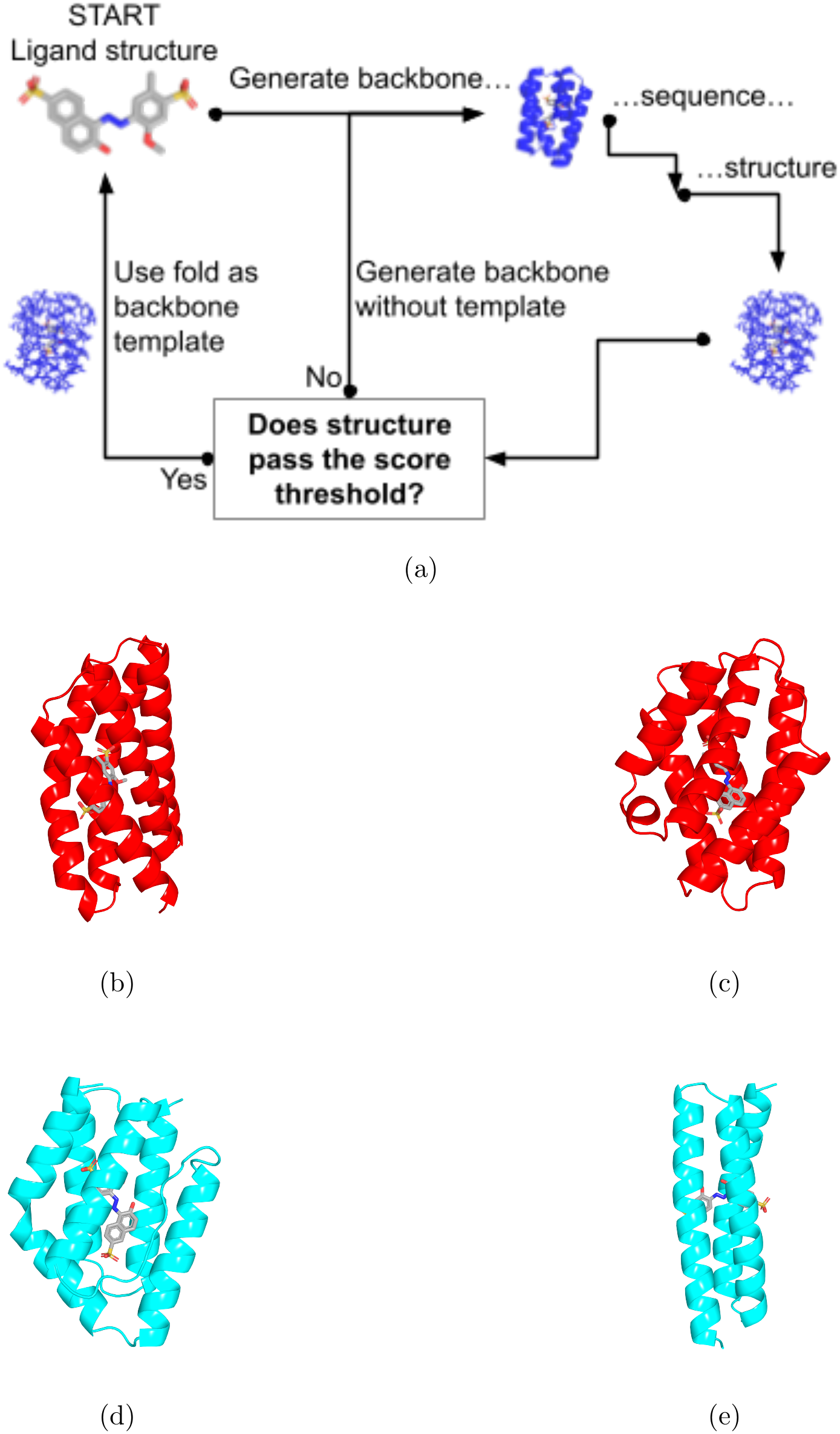
(a) The *de novo* small molecule binder design workflow. (b,c) The median and top-ranked designs from the nonadaptive reference campaign as measured by mean pLDDT. (d,e) The median and top-ranked designs from the IMPRESS campaign.

**Table 1:**
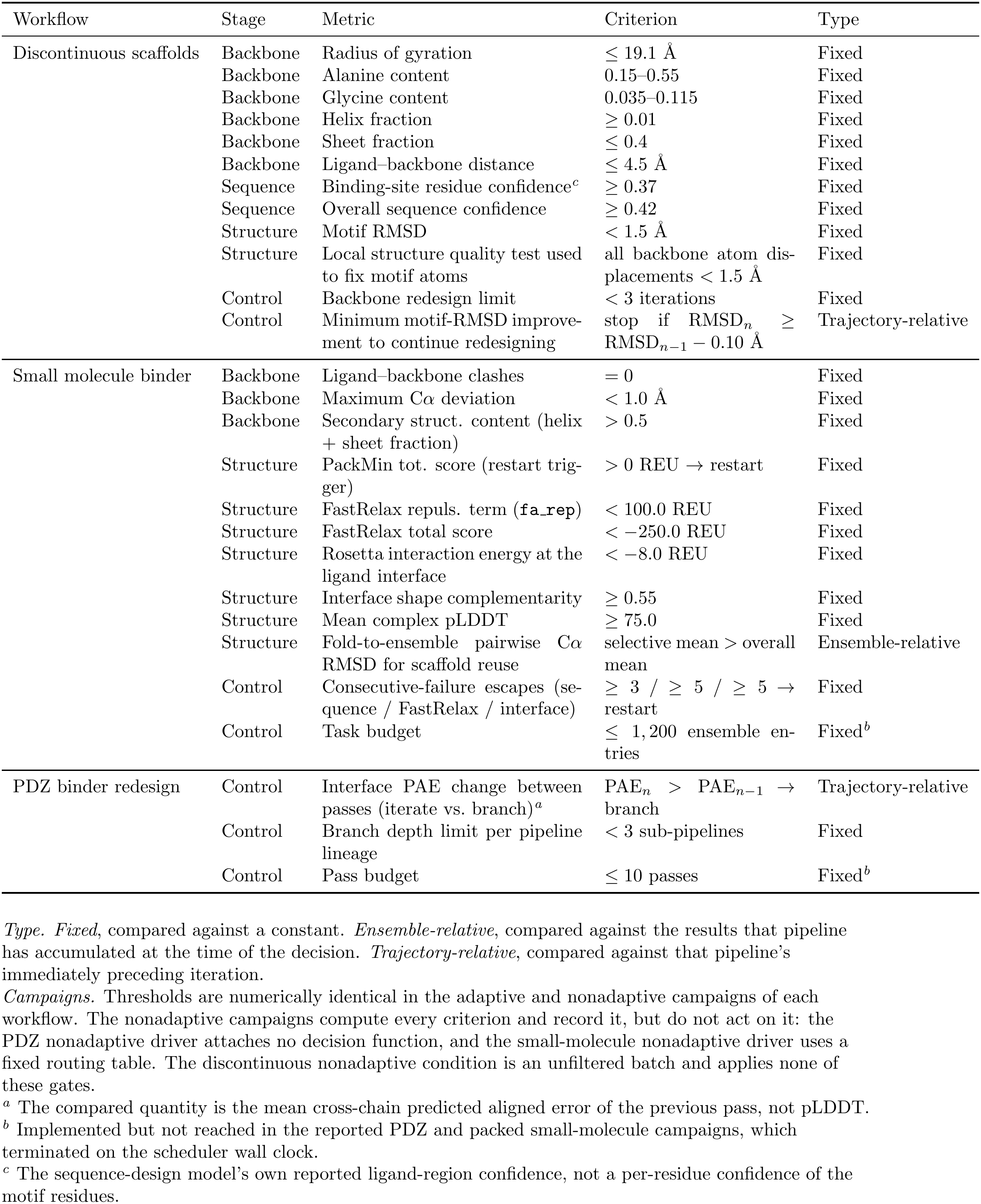
Acceptance gates applied in each workflow.

First, an ensemble of backbone structures containing the binding site is sampled via a diffusion process.^1^ Backbones with suitable gross structure are subject to an iterative sampling scheme in which a sequence is selected and refined based on coarse side chain packing *in situ* on the fixed backbone.^40^ A residue-level structure is predicted for each sequence and is subject to physical scoring, with low-scoring sequences discarded.^41,42^ Among passing sequences, the fold is predicted at atomistic resolution, and the predicted interface between the ligand and binder is ranked on the shape complementarity and predicted conformational stability of the complex.^9,43^

### 2.3 Peptide binder redesign

Conserved sequences and structures in protein-protein binding motifs provide an opportunity for affinity tuning.^27,44^ A bound structure, determined by experiments and comprising a binding protein and a short peptide ligand, is used as a starting point for iterative redesign.

In the peptide binder redesign workflow (Figure 4), a given PDZ domain is redesigned to target a new PDZ binding motif.^26,27^ A sequence is inferred from the input structure containing the modified complex. If the redesigned sequence yields an improvement to the confidence in the predicted fold, then that predicted fold is passed forward for iterative sequence redesign. If the refinement attempt fails, an alternative sequence is resampled from the last iterative step.

**Figure 4:**
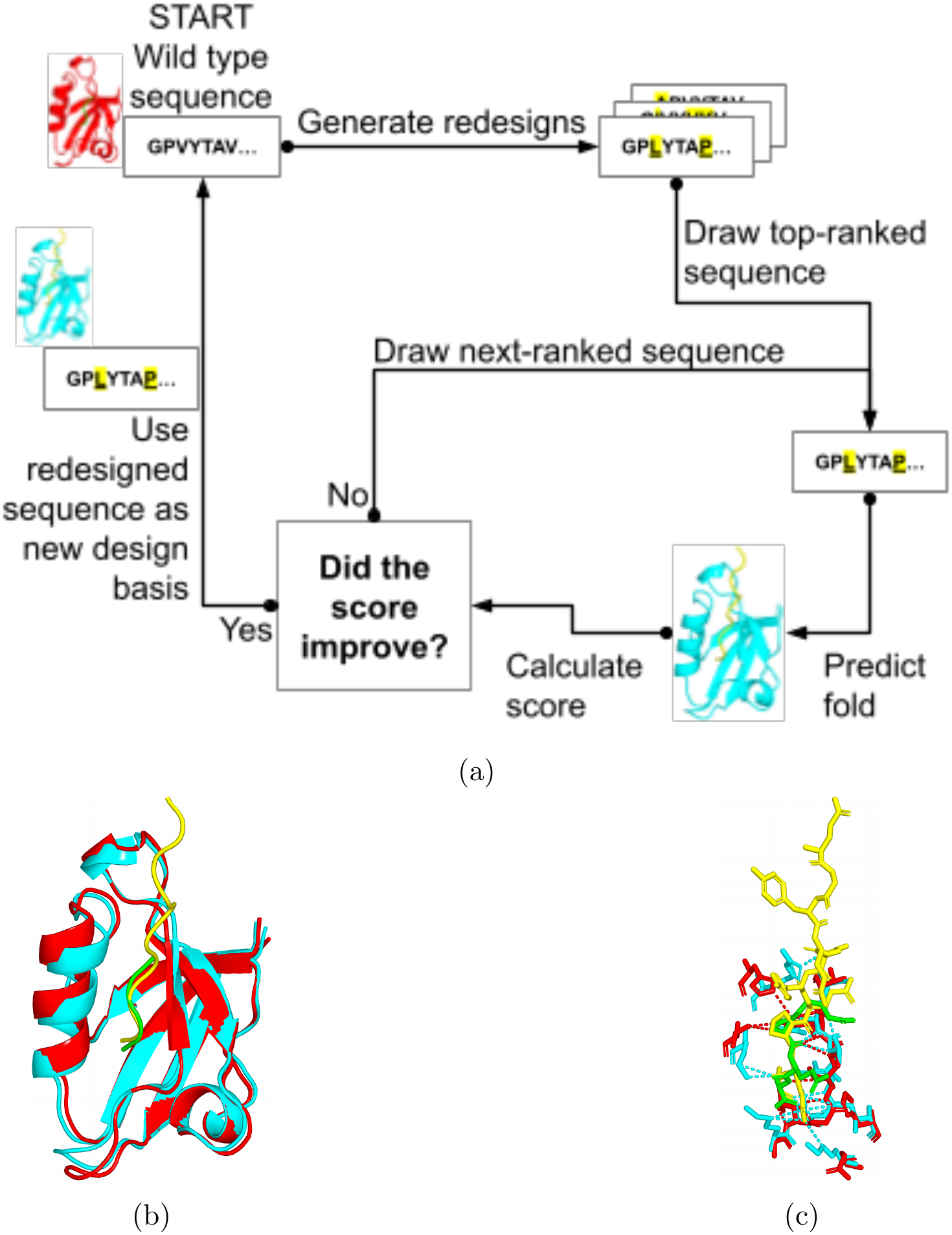
(a) The PDZ binder redesign workflow. (b) Overlay of an example design input (binder in red cartoon, peptide in green ribbon) and a candidate design output (binder in cyan cartoon, peptide in yellow ribbon). (c) Overlay of the interface residues from the input and final designs, showing hydrogen bond (dotted lines) corresponding to each design. The hydrogen bond profile is reworked and expanded in the redesigned protein despite only very minor changes to the secondary structure.

## 3 Methods

IMPRESS is designed to support adaptive protein design workflows that combine AI and simulation tasks on heterogeneous HPC resources while coupling AI to workflow control via the pipeline state.^25^ These workflows are inherently dynamic, as intermediate results continuously influence subsequent computational decisions and determine the workflow’s structure at runtime. Moreover, protein design campaigns often comprise large collections of independent pipelines with diverse and evolving computational requirements.

To efficiently support these workloads, IMPRESS adopts an asynchronous, closed-loop execution model that combines adaptive orchestration with heterogeneous task execution. In the following sections, we present the design principles and architecture of IMPRESS, followed by the implementation details of the framework.

### 3.1 IMPRESS Design

Figure 1 illustrates the design of the IMPRESS framework. Following the original IMPRESS architecture,^22^ the design of the framework follows a building-block approach in which each component is self-isolated, loosely coupled, and independently extensible, allowing individual subsystems to evolve without affecting the remainder of the framework. IMPRESS consists of three core components: (i) the IMPRESS Manager, which performs global orchestration and adaptive decision making (Figure 1, 1–3); (ii) RADICAL-AsyncFlow,^23^ which provides coordination and management of asynchronous graphs (Figure 1, 4); and (iii) the RHAPSODY AI-HPC execution Runtime,^24^ which abstracts the underlying heterogeneous computing resources (Figure 1, 5–6).

**Figure 5:**
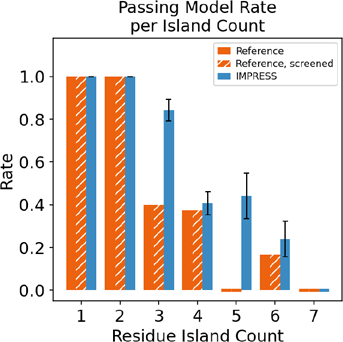
Fraction of input motifs for which at least one predicted structure falls below 1.5 Å RMSD, binned by residue island count, over the benchmark subset of 36 AME motifs folded by both the IMPRESS campaign and nonadaptive reference. Three arms are shown: the unfiltered nonadaptive reference as run; the derived filtered nonadaptive cascade of Section 4.1.1, obtained by applying the Table 1 gates once to the reference’s own recorded criteria; and the adaptive IMPRESS campaign. IMPRESS is truncated to the reference’s 200 folds per motif. Truncation splits taken in campaign order (full initial branch plus 40 of 160 first-redesign folds); motifs whose only passing folds fall in that partial round are scored by the exact probability a random draw of 40 catches one. The 16 single-branch motifs hold only 160 folds and are unaffected, so the truncation is conservative toward the reference. Counts per stratum are given in the text; the filtered and unfiltered reference arms coincide in every stratum, so filtering changes the fold budget spent but not the set of motifs solved. The IMPRESS advantage at 5 islands survives the matched budget, but that stratum holds 4 motifs and is reported as a stratum rather than as a claim; the paired result over the 29 motifs with three or more islands is the quantitative comparison. Error bars on the adaptive arm are the Poisson-binomial standard deviation over which 40 of a partially funded branch’s 160 folds the budget paid for, so they are spread over the truncation and not a confidence interval on the rate; the nonadaptive arms are exact counts over the same motifs and carry no such spread. A rate of zero is drawn as a mark on the axis, to distinguish it from an absent bar.

At the application level, IMPRESS represents scientific workflows as collections of stateful pipelines (Figure 1, 1). Each pipeline consists of a dynamically generated sequence of tasks and maintains its own execution state and intermediate results. Pipelines may encounter adaptive decision points that invoke user-defined functions or LLM-based reasoning components to determine subsequent computational actions.

A decision point is a function over pipeline state, and the framework does not constrain what that function is; the deposited repository contains an example that routes a decision through a language model. Every campaign reported in this work used numerical decision functions only, evaluated against the thresholds of Table 1. No result below depends on a language model.

At each adaptive decision point, IMPRESS first evaluates task outputs against a threshold on an intrinsic metric. Outputs that fail to meet the threshold trigger resampling at the current step. For outputs that pass, an extrinsic metric guides the selection of the next action. The workflow either continues processing with the current results or reparameterizes a subsequent task based on them.

The IMPRESS Manager continuously monitors the collective state of all active pipelines and maintains a global view of their progress (Figure 1, 2). Based on intermediate results and application-specific metrics, the manager dynamically determines whether to stop execution, invoke an adaptive function, and which pipelines or tasks to prioritize. These decisions are materialized through the Pipeline Manager (Figure 1, 3), which constructs and updates the set of computational tasks to reflect the required workflow changes.

The central architectural contribution of the proposed IMPRESS design is the separation of scientific decision-making from asynchronous workflow coordination and heterogeneous task execution. IMPRESS evaluates intermediate scientific results and determines how workflows should evolve, while RADICAL-AsyncFlow coordinates asynchronous task progression and RHAPSODY provides the execution integration for heterogeneous HPC resources. Compared with the previously published approach,^22^ the redesign separates adaptive decision-making from execution and provides a scale-free approach through RHAPSODY without requiring IMPRESS to implement it. Results from completed tasks can guide subsequent work while independent pipelines continue running, without global synchronization.

### 3.2 Implementation

We implemented the IMPRESS framework in Python using RADICAL-AsyncFlow and the RHAPSODY AI-HPC runtime. AsyncFlow is used to express asynchronous task dependencies and workflow execution, while RHAPSODY provides the execution backend for heterogeneous AI and simulation tasks on HPC resources.

IMPRESS introduces two implementation abstractions on top of AsyncFlow: the IMPRESS Manager and the Pipeline. The IMPRESS Manager is responsible for the lifecycle of all active pipelines. It instantiates new pipelines, monitors their execution state, invokes adaptive functions at decision points, and dynamically submits newly generated child pipelines. This event-driven implementation allows adaptive decisions to proceed concurrently with the execution of other pipelines without interrupting ongoing computations.

Each protein design workflow is implemented as a stateful Pipeline object that encapsulates the workflow tasks, execution state, intermediate results, and quality metrics. Rather than representing a static workflow, the pipeline progresses through iterative execution stages and explicitly invokes adaptive decision points after evaluating intermediate results. The adaptive function may terminate the current pipeline, continue execution, or spawn one or more child pipelines that explore alternative design trajectories. Newly created pipelines inherit the relevant execution state from their parent while maintaining independent execution and adaptation. Threshold values used in the production workflows are shown in Table 1.

Pipelines are expressed as asynchronous AsyncFlow graphs of tasks, including both local processing stages (e.g., sequence ranking and FASTA generation) and remote AI/HPC tasks (e.g., ProteinMPNN and AlphaFold).^2,3^ Independent tasks within a pipeline, such as structure predictions for multiple candidate sequences, are submitted concurrently whenever possible, while preserving application-defined execution dependencies.

Task execution is delegated to the RHAPSODY runtime, which schedules heterogeneous AI and simulation tasks through pluggable execution backends. This implementation enables IMPRESS to continuously construct, adapt, and execute large collections of protein design pipelines while efficiently exploiting heterogeneous HPC resources.

### 3.3 Statistical analysis

Every comparison reported in this work pairs an adaptive campaign against a nonadaptive reference built from the same task graph and the same numerical thresholds. The unit of comparison is the design input – an active-site motif or a starting scaffold – since the same input enters both arms. Comparisons are therefore paired by input rather than drawn between pooled sets of designs, which matters because the arms do not contribute equal numbers of designs per input.

For binary outcomes, such as whether a motif was solved, we report counts and the exact (Clopper–Pearson) binomial confidence interval, and test paired outcomes with the exact McNemar test. Where one arm’s successes are a strict superset of the other’s, that test reduces to a one-sided binomial on the discordant pairs, and we say so. For continuous outcomes we report the median or mean paired difference with a bootstrap confidence interval over 10,000 resamples, and test with the Wilcoxon signed-rank test, quoting the rank-biserial correlation as an effect size. All intervals are 95%. Where a test does not reach significance we report the confidence interval alongside it, so that the result bounds the difference that could have gone undetected rather than asserting equality. Tests reported for the same data are not corrected for multiplicity, and we note where that bears on a marginal result.

Each reported campaign is a single execution of a stochastic pipeline; replicate campaigns at this allocation scale were impractical, and we note where a partial replicate exists. The sequence-design and structure-prediction stages were run with fixed seeds (Supporting Information Tables S1–S3), but the generative backbone stage was unseeded, so an individual campaign is not reproducible run-for-run even though its configuration is fully recorded. Pairing by input is what makes the comparisons robust to that variation, and we flag the two cases in which a paired difference could instead reflect it.

## 4 Results

The architectures explored in this paper represent the tip of the iceberg in the sophistication of protein design workflows. As computational architectures grow more sophisticated, they increasingly rely on heterogeneous HPC combining generative ML on GPU with physics pipelines on MPI, but also on external resources including hidden-weights hosted inference models, database APIs for retrieval augmented generation, and large language model (LLM) endpoints. Programs like the National Artificial Intelligence Research Resource^45,46^ (NAIRR), the Genesis Mission,^47^ and the American Science Cloud^48^ foster access to collections that include all of these types of resources, allowing researchers to experiment with sophisticated workflow topologies.

We designed the software, including the example workflows, to target the class of hardware most commonly used by protein design researchers in the academic sphere. Development and testing were performed on HPC allocations provided by the NAIRR Pilot^45,46^ program on the Delta^49^ and Anvil^50^ supercomputers. The NAIRR resources enabled the developers to prove the IMPRESS capabilities on modern heterogeneous high-performance clusters. Many challenges to deployment were systematically addressed in the development of the IMPRESS system and the underlying middleware, including dynamic GPU pinning at the task level and other unforeseen blockers of the asynchronous execution architecture. The NAIRR resources were thus critical to our ability to refine performant architectures capable of accommodating workflows that represent the state of the art.

Production for all three workflows was run on HPC, each against a nonadaptive reference built from the same task graph and the same numerical thresholds. The arms differ in what they do with the measurements rather than in which measurements they take: the PDZ reference attaches no decision function, the small-molecule reference routes tasks from a fixed table, and the discontinuous scaffolds reference is an unfiltered batch that records every criterion without acting on any of them (Table 1). The discontinuous scaffolds workflow was evaluated on the AME benchmark,^6,12^ a curated set of 41 enzyme-ligand binding motifs taken from experimentally determined bound structures, with the catalytic residues and their ligand-bound geometry specified. Table 2 gives the outcome of each comparison alongside the adaptive mechanism responsible for it.

**Table 2:**
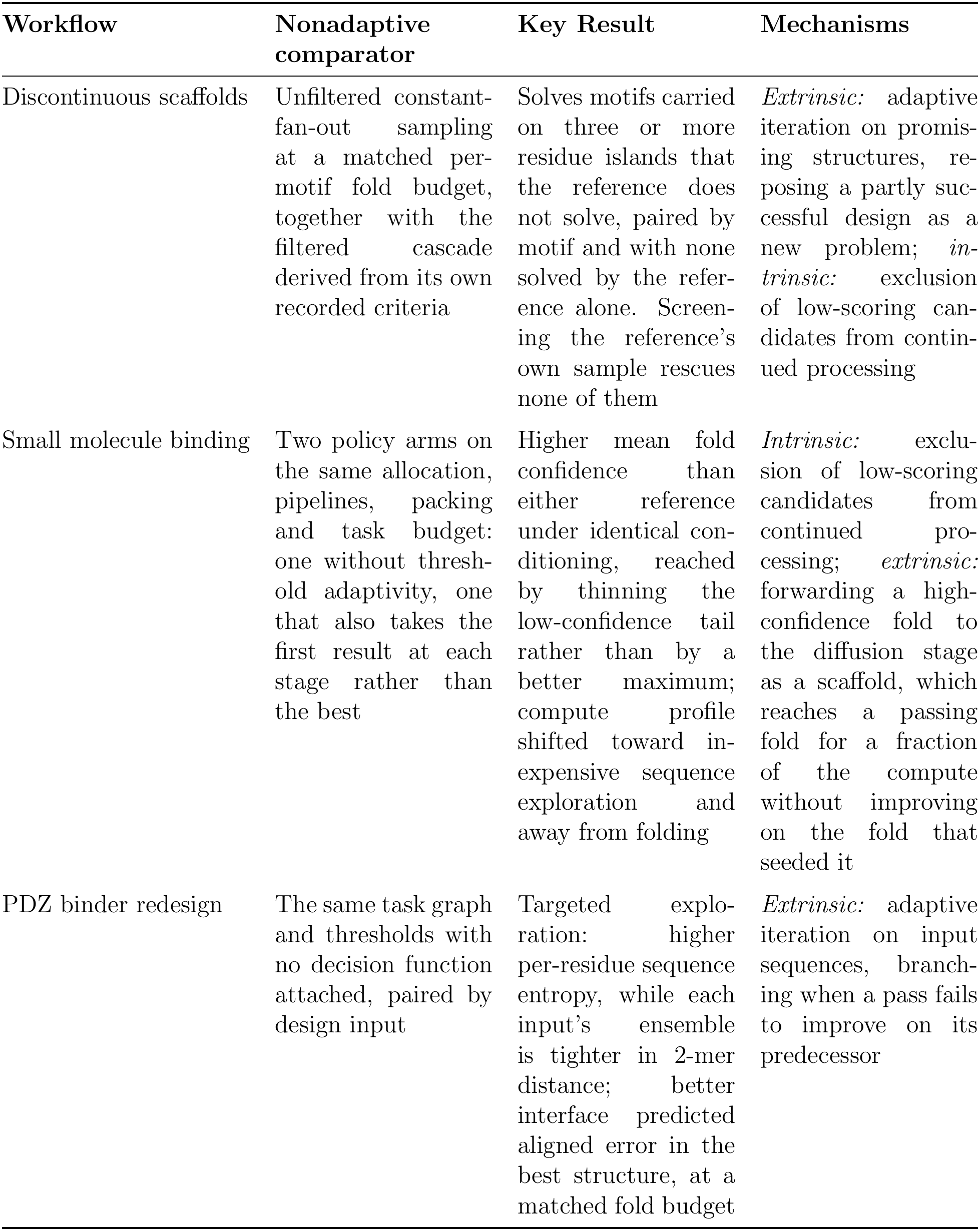
Adaptive workflow outcomes, comparators and enabling mechanisms.

### 4.1 Backbone generation on discontinuous scaffolds

The task pipeline of the discontinuous scaffolds workflow developed here is based on one used to characterize the performance of RFdiffusion on a curated battery of enzyme-ligand poses.^6,12^ The ML models in the pipeline can be used efficiently for batch inference on heterogeneous inputs, so the transformation pipeline can be applied to an entire benchmark library at once.

The AME benchmark library of binding motifs is entered into the pipeline, beginning with backbone generation. Thresholds on secondary structure, alanine and glycine content, and radius of gyration are used to exclude doomed backbones, and sequences are thrown out if residues near the binding site have low confidence scores. The input backbones associated with cancelled sequences are recycled up to a fixed number of times before being discarded.

A nonadaptive reference campaign was run over all 41 AME motifs, closely reproducing the model pass rates reported for RFdiffusion3 on AME.^12^ The reference is an unfiltered batch: it computes and records every criterion of Table 1 but acts on none of them, and its fan-out is constant at 10 backbones, 40 sequences and 200 fold predictions per motif. The comparison is therefore against unfiltered sampling rather than against a filter cascade, and bounds filtering and adaptivity together; Section 4.1.1 separates them. An adaptive campaign targeted a subset of 36 motifs, and results were compared against the matched subset from the reference.

The 36 are the motifs both arms folded, so the reduction is a completion criterion and not a quality filter. None of the five excluded motifs was solved by the reference, so dropping them raises the reference pass rate rather than lowering it. Two of the five are asymmetric cases that cut in opposite directions, and the Supporting Information names all five and states the direction of each. Two of the 36 reached the scheduler wall clock mid-step and are scored here as failures, so an incomplete adaptive run counts against the adaptive arm.

The nonadaptive campaign generated 7,200 fold predictions over 1,440 Chai-1 batches^51^ on the matched subset, and the adaptive campaign 9,716 folds over 1,945 batches, i.e. 200 and 270 fold predictions per motif, respectively. The adaptive arm therefore ran with roughly 1.35 times the per-target fold budget of the reference. To account for this mismatch in the allocations, our post-hoc analysis is limited to the first 200 folds per motif, on the assumption that matching budget on the most expensive single operation gives rough parity in budget.

Figure 5 shows motif solution rates by island count for the unfiltered reference, the derived filtered reference of Section 4.1.1, and the adaptive campaign. Among motifs with 1 or 2 residue islands the three arms are indistinguishable, each solving every motif. The comparison is therefore over the 29 motifs carrying three or more residue islands, paired because the same motif enters both arms. The adaptive workflow solves 17 of the 29 (95% CI 0.389–0.765) against the reference’s 8 (95% CI 0.127–0.472), and its solved set is a strict superset of the reference’s: 8 motifs are solved by both arms, 9 by the adaptive arm alone, none by the reference alone. The exact McNemar test on the 9 discordant motifs reduces to a one-sided binomial, *p* = 0.0020.

Two qualifications bound that result. First, not every discordant motif is evidence of steering, and the data say which are. Two passed on their initial branch, so no adaptive move was made on them. In two others the redesign branch re-entered diffusion from the motif alone, because the parent fold recorded no well-predicted region to carry forward, so that branch is a restart rather than a refinement. Five of the 9 therefore rest on a redesign that inherited structure from the attempt before it, and restricting the test to those five gives *p* = 0.031; the looser restriction, to the 7 on which any redesign branch ran, gives *p* = 0.0078. For the four excluded by the stricter rule the difference is as consistent with sampling variation as with steering, since the reference arm diffused 10 backbones per input against the adaptive arm’s 8 and ran on different hardware with an unseeded generative stage. Second, the budget truncation scores a motif whose only passing folds fall in the partial round by the probability that a random draw of 40 from 160 catches one, so a few reported successes are expected values rather than observations. At 5 islands one success is directly observed (M0732 1xs1, fully funded at 160 folds) and one is imputed at *p* = 0.768 (M0078 1al6); at 3 islands four of the five boundary motifs carry *p ≥* 0.985 and only M0110 1c0p is uncertain at *p* = 0.439; at 4 islands M0349 1e3v carries *p* = 0.25.

The five-island stratum, which earlier drafts of this work reported as the headline, is reported here as a stratum and not as a claim. It contains 4 motifs. The reference solves none of them (95% CI 0.000–0.602), the adaptive workflow solves 2 at its full budget (95% CI 0.068–0.932) and 1 within the matched 200-fold budget on directly observed passes alone. Both of the two fall outside the five discordant motifs identified above: one is the single-branch lineage and the other the bare-motif restart. With that denominator, and on that attribution, the stratum cannot carry a quantitative claim on its own and we do not rest one on it.

That the adaptive campaign solves motifs with three or more residue islands which the reference does not solve at all, under a matched fold budget and with no motif solved by the reference alone, indicates that the adaptive parameterization strategy reaches regions of the design space a fixed pipeline does not, rather than sampling the same regions more cheaply. Further details of motif-level pass rates are shown in the Supporting Information Figure S1.

#### 4.1.1 Filtering alone does not account for the difference

The reference campaign computed every criterion of Table 1 and recorded it without acting on it, so a filtered but nonadaptive cascade over the same sample is recoverable from the retained data and needs no separate campaign. Filtering is a function of quantities the run already measured. We apply the backbone and sequence gates of Table 1 once to the reference arm, in pipeline order, and ask which fold predictions a cascade would have computed and which motifs it would have solved.

Over the reference’s full 41-motif campaign, 333 of its 410 backbones (81.2%) satisfy all six backbone gates, and 1,319 of its 1,640 sequences (80.4%) descend from a surviving backbone and satisfy both sequence gates, so a cascade would have computed 6,595 of the 8,200 fold predictions and saved 19.6% of the most expensive stage. Restricted to the 36 matched motifs the comparison uses elsewhere, the saving is 15.6%: 307 of 360 backbones (85.3%) and 1,215 of 1,440 sequences (84.4%) survive, and the cascade computes 6,075 of 7,200 folds. On that matched subset it solves the same motifs as the unfiltered reference: 4 of 10 at 3 islands, 3 of 8 at 4 islands, none of 4 at 5 islands, 1 of 6 at 6 islands and none of the single 7-island motif, so 8 of 29 pooled over three or more islands, identical in every stratum (Figure 5).

Gating therefore buys economy and not reach. The construction holds the realized sample path fixed, so it describes a cascade that discards candidates without reinvesting the budget it frees, and is in that sense a lower bound on filtering’s cost and an upper bound on its benefit. Reinvestment would not change the conclusion at the hard end of the benchmark: the freed budget corresponds to 1.24 times as many folds, while all four five-island motifs have no passing fold anywhere in the reference’s 200, so a larger nonadaptive sample has nothing to recover. The comparison that isolates adaptivity at fixed gates and thresholds is the small-molecule experiment of Section 4.2, whose reference applies the same gates from a fixed routing table.

### 4.2 *De novo* small-molecule binders

The small-molecule binder workflow comprises the usual set of diffusion pipeline design transformations – backbone generation, sequence prediction, and fold prediction – plus analysis and adaptive control after each transformation. Each design run is realized as a random sequence of transformation tasks based on control flow rules. Results in the three design modalities – backbone, sequence, and folded structure – accumulate as a trio of ensembles of design candidates.

The workflow is initialized with a user-supplied target ligand structure that feeds into the first round of backbone diffusion. Subsequently, runtime conditions determine whether designs progress forward through the backbone-sequence-fold pipeline or cycle backward to resample at a previous step. The adaptive controls induce a random ordering of transformation tasks. The proportions of the realized task sequence can be shaped via tuning of a small number of physically interpretable parameters. These parameters control the pace at which the workflow explores the design space.

Compute duration is bounded jointly by a per-pipeline task budget and by the scheduler wall clock. Which of the two binds depends on how productive each pipeline is: in the packed configuration described below no pipeline reached its task budget within the four-hour allocation, whereas in the unpacked configuration, where each pipeline had a GPU to itself, 11 of 32 pipelines exhausted the budget before the wall clock. Reported throughput for that arm is therefore a lower bound on what the allocation could have delivered.

All generated backbones are evaluated for secondary structure content and for similarity to the ensemble of backbones generated during the run measured as pairwise RMSD. Backbone prediction is rerun until one with enough secondary structure is produced, and suitable backbones are processed forward to sequence prediction. The evaluation of each sequence continues with an inexpensive structure prediction with Rosetta,^41^ providing a basis for gating of the interaction energy at the ligand interface. Fold predictions at atomistic resolution are made with Boltz-2.^9^ Folds are scored and gated on conformation confidence, as Boltz-2 mean complex pLDDT over all tokens in the bound complex, and interface energy.^3^ Passing folds are compared with the ensemble via pairwise RMSD; folds similar to past high scorers are passed as scaffolds for backbone prediction. Adaptive gating followed the intrinsic and extrinsic scheme described in Methods, with the thresholds of Table 1.

Experiments were run to evaluate the performance-tuned adaptive workflow and compare it with two allocation-matched nonadaptive reference workflows, the first of which forgoes threshold adaptivity (pass-through) and the second of which also forgoes ranking of candidates per stage instead using the first result produced (naive). The three campaigns are matched on every recorded run condition: each occupied one node and four A40 GPUs, ran the same 32 pipelines packed eight to a GPU on the same pipeline-to-device map, used the same 60-second staggered start and the same budget of 1,200 ensemble entries per pipeline, billed the same allocation, and ended on the four-hour scheduler wall clock rather than on the task budget. The decision policy is the only field that differs, so the comparison below is a controlled ablation. A fourth run repeated the adaptive arm on a different node under the same conditions and is reported as a consistency check rather than as a replicate arm: it returned a mean fold confidence of 89.3 against the primary adaptive run’s 89.4 and a median of 92.7 against 93.2, from a smaller ensemble. We note that the packing comparison later in this section is not matched in this way.

Every fold computed in every arm is retained and scored, including those the gates rejected: across the three arms 2,789 designs entered the campaign comparison and 2,678 of them returned a fold prediction, of which 1,548 satisfy the design-stage acceptance criteria. The distributions of Figure 6 are drawn over the latter population, in which all three arms are conditioned identically on the FastRelax and interface criteria. Conditioning is necessary here because the adaptive policy halts a design that fails the FastRelax criterion before the interface calculation runs, so that design never enters the adaptive arm’s ensemble while a nonadaptive arm carries its failures forward; comparing the ensembles as produced would compare a screened population against an unscreened one. The fold confidence criterion is excluded from the conditioning, since it is the quantity being compared and screening on it would presuppose the result. The comparison is insensitive to the choice: over every computed fold the adaptive arm’s mean complex pLDDT exceeds the pass-through reference’s by 0.061 (0.894 against 0.833), and under identical conditioning by 0.052 (0.894 against 0.841), the reference gaining slightly and the adaptive arm not at all, since its ensemble is already screened.

**Figure 6:**
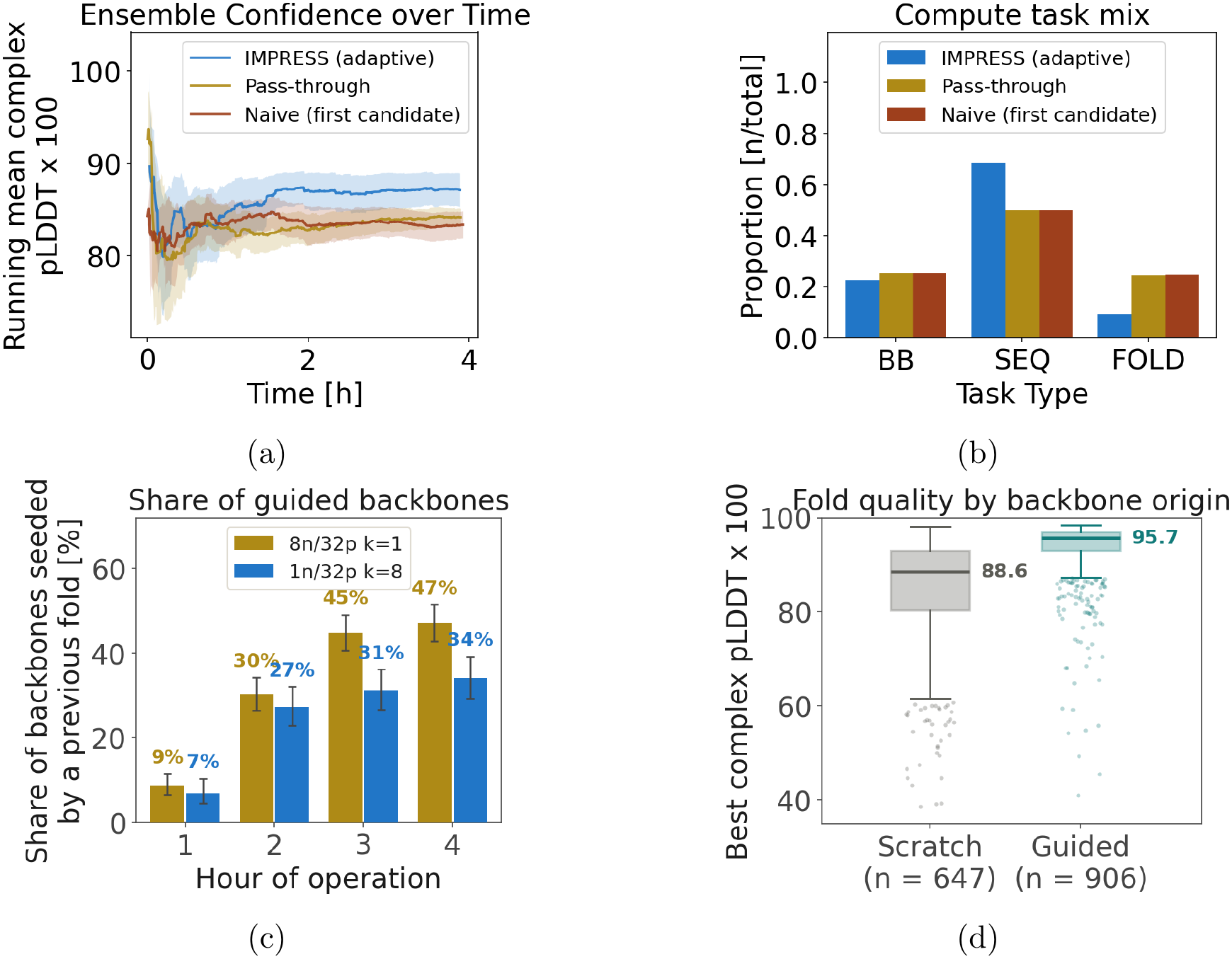
Adaptive moves in the small-molecule binder workflow and their effect on the realized campaign. Panels (a) and (b) compare the adaptive campaign with its nonadaptive references; panels (c) and (d) characterize the scaffold-reuse move within the adaptive campaign.(a) Running average mean Boltz-2 complex pLDDT in IMPRESS vs. nonadaptive reference. IMPRESS samples higher and more broadly, and converges later than reference. All three arms are conditioned identically on the FastRelax and interface criteria, which the adaptive policy applies during the run and the references do not; fold confidence, the quantity plotted, is excluded from that conditioning. The ordering is unchanged over every computed fold. (b) Compute task mix in the IMPRESS and reference campaigns. IMPRESS performs more (inexpensive) sequence redesigns per (expensive) fold inference than does reference. Counted over task directories rather than compiled designs, so no conditioning applies. (c) Share of backbones seeded by a previous fold at each hour of operation in two packing conditions, 1 pipeline per GPU and 8 pipelines per GPU. Whiskers are Wilson intervals on the hourly proportion. The fourth hour of the 1-pipeline-per-GPU run excludes 11 of its 32 pipelines, which stopped on a task budget of 300 at minute 202. (d) Distribution of structures over best complex pLDDT among scaffold-guided backbones vs. *de novo*. Seeded designs are distributed more tightly at the high end, with a lower quartile of 93.0 against 80.4, but this compares populations selected on different criteria rather than treatments: the move fires only on a fold that has already passed, so a seeded design begins from a known-good parent. Measured against its own parent fold, a seeded design is no better (median +0.010 in complex pLDDT over 906 pairs, 50.3% improving, Wilcoxon *p* = 0.86). The benefit is economic: 85.2% of seeded episodes reach a passing fold against 19.5% begun from scratch, or 8.3 tool invocations per passing fold against 33.1.

Figure 6 shows the time evolution of the ensemble mean pLDDT generated by IMPRESS and nonadaptive reference. The adaptive ensemble reaches a higher quality than either reference and does so by thinning its low-confidence tail rather than by extending its upper end. Under identical conditioning, 8.1% of adaptive folds fall below the workflow’s own 0.75 confidence criterion against 19.3% and 19.9% for the pass-through and naive references, and the medians are 0.932 against 0.881 and 0.878 (Mann–Whitney *p* = 4.6 *×* 10*^−^*^22^ against the pass-through reference, rank-biserial +0.34; these arms are not paired, since each generates its own backbones). The best single design differs little between arms – 0.983 against 0.979 and 0.977 – so the result is a shift of the distribution and not a better maximum, which is the comparison that matters when the arms hold ensembles of different size.

Because mean complex pLDDT is averaged over all tokens in the complex, it is a weak proxy for the quality of the binding interface specifically, which is the quantity of interest and which the PDZ workflow of Section 4.3 measures directly as interface PAE. Two interface-local Boltz-2 scores are recorded for every one of those 2,678 folded designs and agree with the pLDDT ranking under identical conditioning: interface predicted TM-score is 0.848 for the adaptive arm against 0.792 and 0.789 for the pass-through and naive references, and interface pLDDT 0.826 against 0.743 and 0.736. The three arms did not produce equal numbers of designs within the allocation – 650 for the adaptive arm against 1,056 and 1,083 for the references – which is itself the task-mix result below. All distribution comparisons are therefore made at matched sample size, every distributional statement is read as a difference in shape rather than in count, and no claim is made from the maximum of an ensemble. The IMPRESS workflow drives proportionally more sequence tasks compared with reference, indicating a shift in computational workload away from expensive folding and toward cheap sequence exploration.

The second adaptive move in this workflow forwards a high-confidence fold to the diffusion stage as a scaffold, and its effect is economic rather than qualitative. The share of backbones seeded from the campaign’s own folds rises over the run as the ensemble accumulates, from 8.8% in the first hour to 47.1% in the fourth (Figure 6c), so the feedback loop does engage. Seeded designs are distributed more tightly at the high end of the quality range than designs begun from scratch (Figure 6d), with a lower quartile of 93.0 against 80.4, but that gap is a consequence of which designs are eligible for seeding rather than evidence that seeding improves a design: the move fires only on a fold that has already passed, so a seeded design begins from a known-good parent. Compared against its own seeding parent, a seeded design is not better – the median change in complex pLDDT over 906 such pairs is +0.010, with 50.3% improving (Wilcoxon *p* = 0.86), and the change in interface predicted TM-score is indistinguishable from zero (*p* = 0.44) – and the gain by parent quality tercile runs +1.46, +0.03, *−*0.17 from low to high, the signature of regression toward the mean rather than of an effect. What seeding buys is throughput: 85.2% of seeded design episodes reach a passing fold against 19.5% of those begun from scratch, so a passing fold costs 8.3 tool invocations by the seeded route against 33.1 by the other, a four-fold reduction. This is the same distinction the active-site benchmark draws in Section 4.1.1 between reaching a design and reaching it cheaply, and here the adaptive move buys the latter.

Performance tuning was achieved via experiments to assess the performance potential of a scheme to pack more pipelines into a fixed allocation. In the unpacked scheme, 32 pipelines were simultaneously deployed with one GPU dedicated per pipeline (8 nodes). In the packed scheme, the same 32 pipelines were started at 60-second intervals and packed 8 to a GPU across 4 GPUs (1 node), delaying GPU allocation for later pipelines and allowing hardware availability to catch up with demand. A brief analysis of the packing exercise is shown in the Supporting Information Figure S2.

Both arms ran the same adaptive workflow, confirmed from their telemetry, which records 9,162 adaptive decision calls in the packed arm and 13,074 in the unpacked arm. They are not, however, matched in allocation or in task budget: the packed arm occupied 1 node and 4 GPUs against the unpacked arm’s 8 nodes and 32 GPUs for the same four-hour span, and was allowed 1,200 ensemble entries per pipeline against the unpacked arm’s 300. Within this pair, placement density and allocation size cannot be separated, since packing 32 pipelines eight to a GPU is what reduces the allocation to one node. A third condition bounds the allocation contribution directly, and is reported in Supporting Information Figure S2: holding placement at one pipeline per GPU and scaling the allocation fourfold, from 8 GPUs on two nodes to 32 GPUs on eight, moves mean GPU occupancy from 11.4% to 12.2%. Allocation size therefore accounts for under a percentage point of the difference, and the rise to 75.8% is attributable to placement density. The task-budget difference binds only in the fourth hour, where 11 of the unpacked arm’s 32 pipelines stop on it, so all comparisons below are drawn inside the first 200 minutes. Results are summarized in Figure 7.

**Figure 7:**
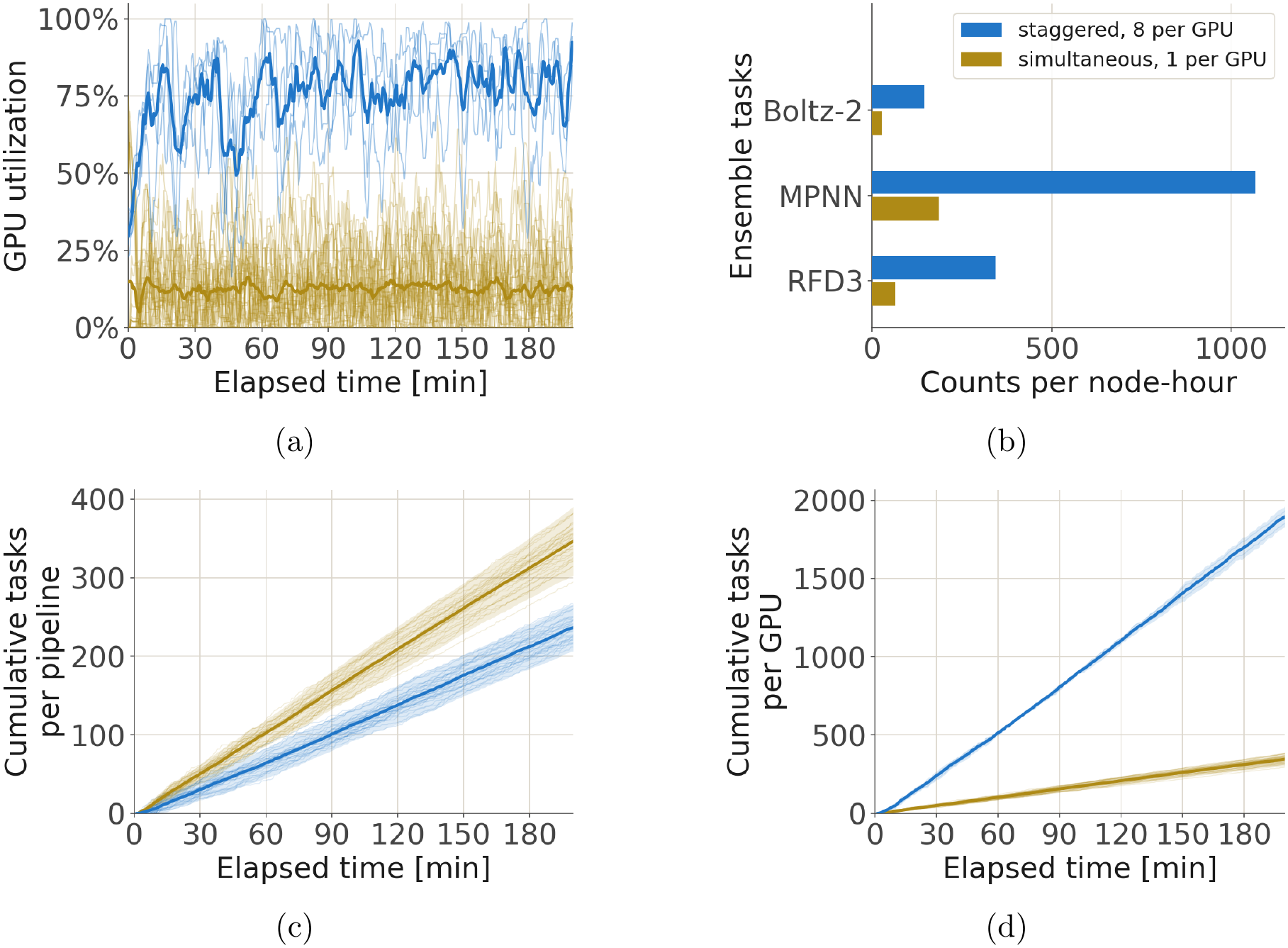
Efficiency and scale of a tuned pipeline. Two four-hour jobs of 32 pipelines each were run, differing only on packing strategy. Ochre: naive packing – 32 simultaneous pipelines, 1 per GPU across eight nodes. Blue: staggered start – 32 pipelines launched in eight groups of four at 60-second intervals, 8 to a GPU. a Built-in telemetry shows GPU utilization in each experiment. Dark colored lines show a 4-minute moving average. Packing fills the nodes: mean utilization 75.8 % against 12.2 % over the full four hours, with the packed node’s four GPUs tightly matched at 74.2–76.7 % where the unpacked run’s 32 range 9.7–15.6 %. The packed run ramps over roughly its first hour as the pipelines fill – 60% over minutes 0–15, 69% over 15–60 – and is flat at about 79% from there to minute 200; the unpacked run is flat throughout. b Task completions per node-hour, counted from minute 10 to each run’s wall clock. Holding pipeline count fixed and varying only the packing gives 5.5× the RFD3 backbones (344.0 vs 63.0), 5.7× the LigandMPNN sequences (1,068 vs 186) and 5.1× the Boltz-2 folds (144.1 vs 28.0), and cuts the cost of a passing fold 5.1×, from 0.159 to 0.031 A40 GPU-hours. c Cumulative tasks per pipeline and d per-GPU. Faint lines are one pipeline (left) or one device (right); heavy lines are the run’s mean and bands are ±2 standard deviations. The unpacked pipelines achieve higher throughput because pipelines never wait for a GPU. Per-GPU ranking inverts that, showing that the packed run returns 511 tasks per GPU-hour against 103 in the naive case. Packing also makes devices more uniform, because each is multiplexing eight independent pipelines whose variation averages out. Because the unpacked run gave each pipeline its own GPU, its ochre curve is the same data in both bottom panels; only the blue curve is redivided.

Packing raised mean GPU utilization from 12.2% to 75.8% and reduced the cost of a passing fold 5.1-fold, from 0.159 to 0.031 A40 GPU-hours. The two statements describe the same effect from different ends: in the same four hours the packed arm consumed 16.0 GPU-hours and returned 512 passing folds, while the unpacked arm consumed 127.9 GPU-hours – eight times as many – and returned 804. In absolute terms the unpacked arm therefore delivered more work, 1,930 backbones and 873 folds against 1,319 and 558, and did so despite 11 of its pipelines stopping on the task budget. The gain from packing is thus an eight-fold reduction in resource cost for roughly two-thirds of the absolute throughput: an efficiency-for-latency trade at fixed pipeline count rather than a throughput win.

Figure 7c shows the per-pipeline side of that trade, in which pipelines sharing a GPU complete fewer tasks each because a pipeline that owns a device never waits for it. For a campaign of independent pipelines, where only aggregate cost per result matters, the trade is worth making; for campaigns over dependent or heterogeneous pipelines, where a slow pipeline delays its successors, the reduction in per-pipeline throughput may matter more than the saving.

Pipeline packing resulted in improvements to GPU utilization (Figure 7a) and throughput per task (Figure 7b) and per GPU (Figure 7d). The per-node-hour and per-GPU figures of Figures 7b and 7d are normalized rates, and the factors of roughly five that they show should be read against the absolute totals above rather than as absolute gains.

### 4.3 Peptide binder redesign

Given a set of candidate binders posed in bound complex with their target ligands, a genetic algorithm is developed to explore the binder sequence redesign space.^26,52^ A set of 53 candidate binders was curated from the PDB,^53^ and the sequence iteration workflow was implemented in IMPRESS. In the workflow, sequences associated with high-quality predicted structures are iterated, with a new sequence generated using the predicted fold. If the iterated sequence results in an improvement, iteration continues. Otherwise, if the predicted structure scores low, a branch pipeline is created and a fresh sequence is predicted for the given backbone. This process allows the sequence features of good designs to pass to genetic descendants, with family trees punctuated by random re-rolls.

The results of redesign campaigns using the adaptive workflow and a nonadaptive reference workflow are shown in Figure 8.

**Figure 8:**
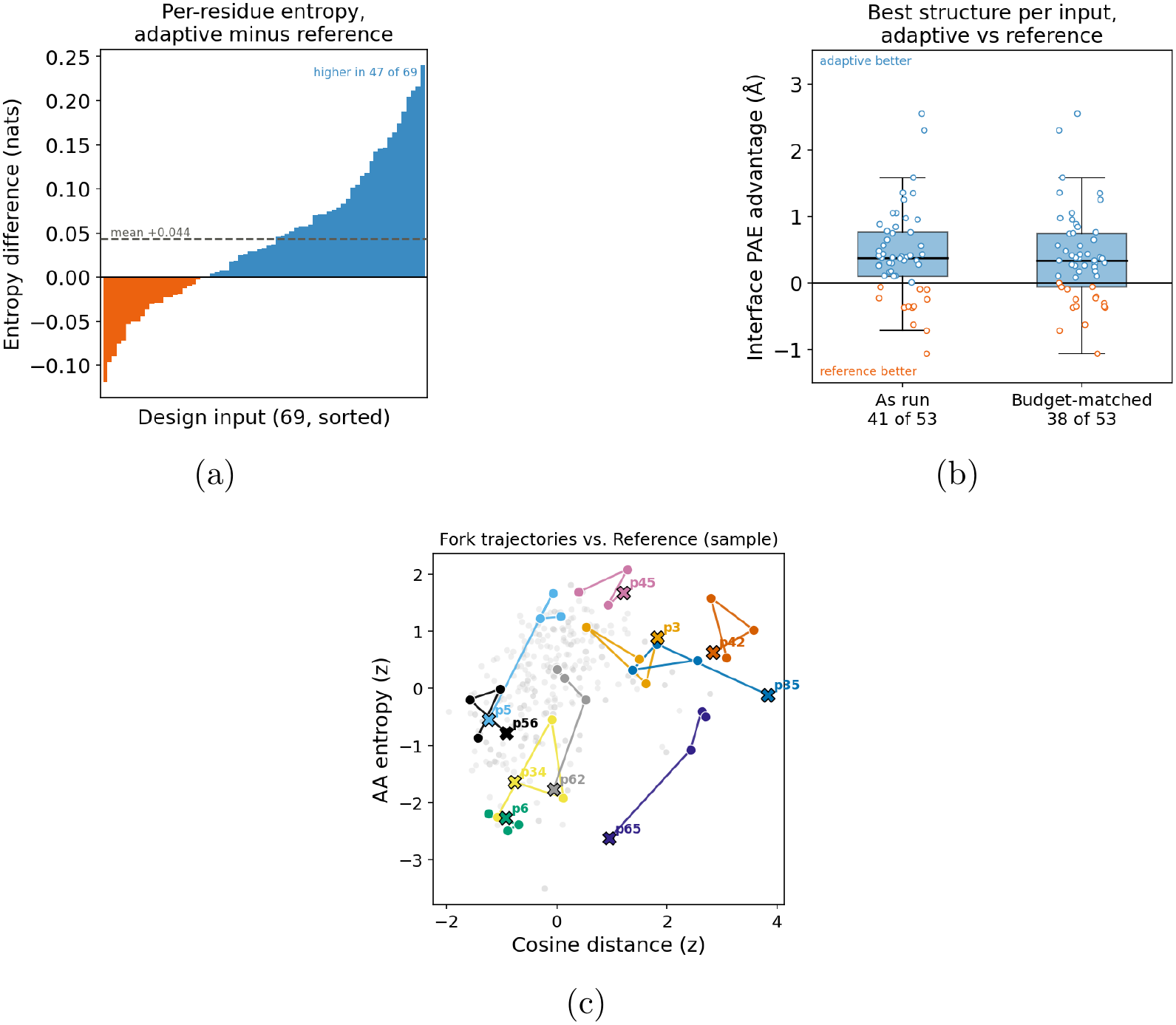
Sequence-space and interface behaviour of the adaptive PDZ redesign campaign. Arms are compared input by input, since the same starting scaffold enters both and they contribute unequal numbers of designs to each: a median of 4 sequences against 1, and 12 folds against 8. (a) Difference in mean per-residue Shannon entropy, adaptive minus reference, for each of the 53 paired inputs, sorted. Entropy is higher in the adaptive arm for 36 of them, a mean increase of 0.043 (95% CI 0.023–0.064; Wilcoxon *W* = 326, *p* = 5.6*×*10*^−^*^4^, rank-biserial +0.54). (b) Advantage of the adaptive arm’s best structure over the matched reference’s, in interface predicted aligned error, for each of the 53 paired inputs. Positive is better. The left group is the comparison as run; in the right group the adaptive arm is truncated to the reference’s fold count for that input, so the two arms spend the same budget. The adaptive arm leads for 41 of 53 as run (77.4%) and 38 of 53 under the matched budget (71.7%, 95% CI 57.7–83.2%; Wilcoxon *p* = 1.2 *×* 10*^−^*^4^), so the result does not depend on the budget gap. Boxes are the interquartile range with the median; whiskers are the 1.5 IQR range and points are individual inputs. (c) A projection of the sequence ensembles onto a per-sequence measure (amino acid entropy) and an inter-sequence measure (mean 2-mer cosine distance), both normalized. The IMPRESS design trajectories typically explore a sequence space that is near the reference input, showing that the redesigns are targeted rather than completely random. Quantitatively, the mean pairwise 2-mer cosine distance within an input’s ensemble is lower in the adaptive arm by 0.0042, but over these 53 inputs the difference is not resolved (95% CI *−*0.0101 to +0.0017; Wilcoxon *p* = 0.23); the interval bounds any ensemble-level difference at 0.010 in magnitude. The targeting visible here is therefore a qualitative feature of the trajectories, and the quantitative sequence-space result is the per-position entropy of panel (a).

The clearest signal is targeted exploration. The units of comparison are matched design inputs – the same starting scaffold enters both arms – so the arms are compared one input at a time rather than as pooled sets of sequences, each reduced to one value per input before testing. On that basis the adaptive workflow explores more widely within each input. Per-residue Shannon entropy is higher in the adaptive arm for 36 of the 53 paired inputs (Figure 8a), a mean paired increase of 0.043 (95% CI 0.023 to 0.064; Wilcoxon signed-rank *W* = 326, *p* = 5.6 *×* 10*^−^*^4^, rank-biserial *r* = 0.54). The corresponding ensemble-level measure does not resolve: the mean pairwise 2-mer cosine distance between an input’s sequences is lower in the adaptive arm by 0.0042, but the interval spans zero (95% CI *−*0.0101 to +0.0017; Wilcoxon *p* = 0.23), which bounds any difference in ensemble spread at 0.010 rather than establishing one. We therefore claim the per-position result and not an ensemble-level one: the adaptive pipelines sample a broader set of residues at each position, the behavior expected of a genetic algorithm that inherits sequence features along a lineage and re-rolls only on failure, while whether they also hold each input’s ensemble closer together is not settled at this sample size. Exploration is targeted rather than random, as the trajectories of Figure 8c show.

Design quality was assessed on the Boltz-2 mean cross-chain predicted aligned error of each input’s best fold, in an input-matched analysis over the same 53 candidates. Matching each input’s budget to the smaller of the two arms, the best adaptive design outscores its matched reference for 38 of the 53 inputs, a win rate of 71.7% (Figure 8b; 95% exact binomial CI 57.7% to 83.2%; Wilcoxon signed-rank *p* = 1.2 *×* 10*^−^*^4^, rank-biserial *r* = 0.61). The comparison as run, in which the adaptive arm held a median of 12 folds per input against the reference’s 8, gives 41 of 53 (77.4%, *p* = 1.9 *×* 10*^−^*^5^), and rarefying instead of truncating gives 39 of 53 (73.6%, *p* = 9.2 *×* 10*^−^*^4^). We quote the budget-matched figure as primary because the as-run comparison is confounded by that budget difference, and report all three because the conclusion does not depend on which is chosen.

Such subtle steering of pipeline dynamics may produce a useful effect in campaigns where promising design neighborhoods are identified. Such subtle steering of pipeline dynamics may produce a useful effect in campaigns where promising design neighborhoods are identified. An analysis of the self-similarity of sequences generated by the adaptive pipeline is shown in Supporting Information Figure S3.

## 5 Discussion

### 5.1 Tuning design ensemble quality at runtime

These results support the premise that the quality of a design ensemble can be tuned by adapting pipeline parameterization to an objective. This performance improvement can only be realized over a large sample of computational experiments such as is common practice in programmatic investigations of molecular docking and binding. ^4^ The use cases presented above span a range of tasks in the practice of protein design in the era of AlphaFold and RFDiffusion.^1,3^ The tasks themselves span a range of compute requirements (CPU, GPU, time). This allows us to characterize the compute demands of each stage of a protein design campaign, and provides a platform for ablation testing to determine the cost of a unit of design quality.

### 5.2 AI-HPC Coupling and the role of the workflow manager

AI and HPC paradigmatically represent the software/hardware decomposition in contemporary protein design. Atom-scale particle-based molecular models retain the unique capacity for producing interpretable free energy surfaces in macromolecular systems.^54^ Inference engines like AlphaFold and RFDiffusion cannot provide dynamical information.^55^ However, their sampling behavior is readily modulated via generalized parameters like temperature, and thus may be practically applied to generate a conditionally converged conformational ensemble that is arbitrarily more or less constrained (or erroneous) than one produced by a carefully tuned parameter set like a force field. Thus ML models may be used to generate a maximally diverse pool of candidate designs – targeting the spatial side of the ergodic hypothesis – whose plausibility is gauged by model-specific ‘gray box’ measures. The *in silico* nature of *plausibility* in this context (ML-native metrics e.g. scores in AF, Rosetta, ESM^3,41,56^) allows error to creep in as a result of biasing to the input data (i.e. the model fails to generalize over the physics, read: PDB-trained models make PDB-looking designs^14,57^).

In multi-step workflows (e.g., backbone-sequence-fold), model fine-tuning can be used for strategic improvement in the pipeline, a methodological shift fueled by flexible technologies like diffusion and transformers.^1,56^ In response, modern middleware frameworks seek to accommodate machinery both for performant task allocation and for management of training loops. This is achieved by situating inference tasks and online training tasks beneath a common control plane, providing a unified basis for fine control of ML workflow behavior (i.e., the character of the designed ensemble in terms of its analytics).

AI-HPC coupled architectures combine black-box inference with adaptive control flow to determine task scheduling at runtime on post-exascale systems.^20,21,25^ The degree or nature of the coupling is mediated by the context shared between the HPC program and the ML model, defined as the collection of mutable variables accessible to the control plane. This enables a variety of AI-coupled HPC motifs to be realized within a single framework, requiring only that the workflow state be designed to support the desired coupling paradigm.

### 5.3 Limitations of ML model self-reported metrics and multifactorial adaptive campaigning

The metrics used to steer these workflows are also typically the metrics used to evaluate them, and this circularity bounds what the results can establish.^15,55^ Further, the adaptive campaigns typically combine multiple metrics and mechanisms for candidate gating and downstream task parameterization. Thus these results do not decisively prove that a particular metric corresponds to an improvement in the objective quality of a protein. Rather, they provide evidence that metrics can be arbitrarily targeted for refinement via instrumented adaptivity, a fundamentally separate concern from the question of whether a particular *in silico* value corresponds to real-world quality.

## 6 Conclusions

The design of generative protein design engines requires iteration over their architecture and implementation. Often multiple ML models or parameterization strategies are considered when planning a particular design task. New modelling tools are published at a prodigious pace, bringing performance improvements or new capabilities. A performance improvement may drive the workflow designer to replace a particular inference model, but a change in capability can drive a revision to the architecture, and the iterative process continues.

IMPRESS is a high-performance workflow composition framework designed to enhance the effectiveness of iterative design in the current environment of fast-paced developments in ML for protein design. The building block approach that IMPRESS takes to the design of workflows supports flexible composition of workflows over the task compute space. This composable machinery is projected onto an asynchronous execution layer that provides high performance in pipeline throughput without the need to re-engineer performance for every revision to the workflow – the user composes the workflow, performance comes for free.

In this work we implement a survey of AI-driven-HPC protein design workflows using design-analyze-adapt cycles to assess the performance of the IMPRESS engine. Our experiments demonstrate that adaptive strategies to improve the protein design ensemble can be deployed through IMPRESS effectively and efficiently, using a variety of adaptive strategies and task tools.

The current era of agentic and autonomous systems has changed workflows design, and new task tools and workflow topologies have proliferated. Modular orchestration systems like IMPRESS provide a means of implementing performant systems for protein design into long-lived tools that are readily refined in response to the rapidly changing AI-enabled computing landscape.

## Acknowledgement

Development of the IMPRESS software was made possible by NSF 2438557 (NAIRR Pilot: IMPRESS: Integrated Machine-learning for PRotein Structures at Scale) and allocation NAIRR 240405.

This research used the Delta advanced computing and data resource which is supported by the National Science Foundation (award OAC 2005572) and the State of Illinois. Delta is a joint effort of the University of Illinois Urbana-Champaign and its National Center for Supercomputing Applications. This research also used the Anvil advanced computing and data resource which is supported by the National Science Foundation (award OAC 2005632). Anvil is a joint effort of Purdue University and its Rosen Center for Advanced Computing.

M.H. was supported by a postdoctoral fellowship from the Rutgers Artificial Intelligence and Data Science Collaboratory (RAD).

Generative AI was used in the production of the software presented in this work and in the editing of the manuscript text. The authors directed, reviewed and take full responsibility for all of it. No result reported here depends on a language model at runtime; all decision functions in the reported campaigns are numerical, as described above.

## 7 Data and Software Availability

The IMPRESS software is freely available under the MIT license at https://github.com/ radical-collaboration/IMPRESS. Workflow definitions for the three use cases as well as the benchmark targets are included in the repository.

## Supporting Information

### Discontinuous scaffolds benchmark

**Figure S1:**
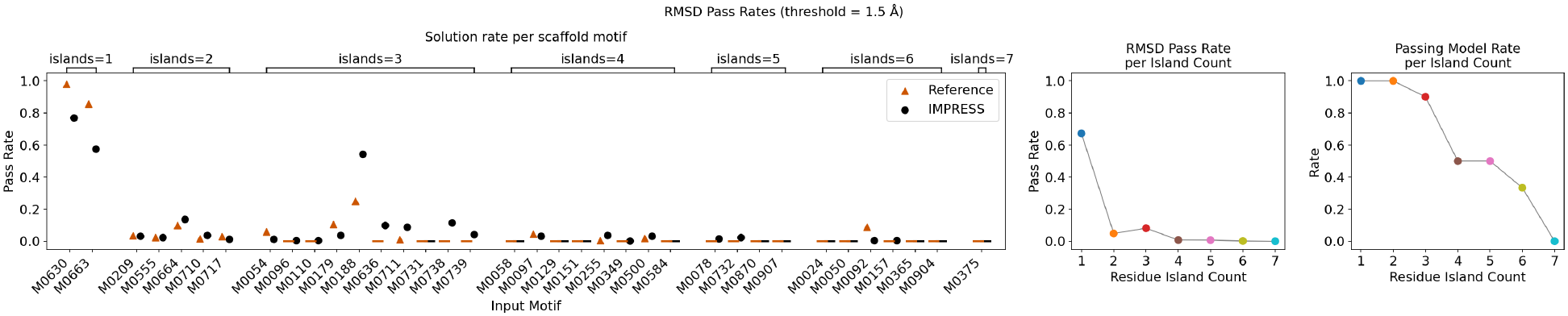
RMSD Pass Rates per motif from the benchmark set. The per-motif pass rates reflect the proportion of candidate solutions from each workflow that produce a solution acceptable to the 1.5-Ang motif RMSD criterion.

The adaptive and nonadaptive campaigns are compared over the 36 motifs both arms folded, out of the 41 in the AME benchmark. The rule is completion, not quality: a motif enters the comparison if the adaptive campaign produced fold predictions for it. The five excluded motifs are M0040 13pk, M0093 1dqa, M0315 1ey3, M0552 1fgh and M0674 1uf7. None was solved by the reference, so excluding them raises the reference pass rate rather than lowering it.

Two of the five are asymmetric, and they cut in opposite directions. M0674 1uf7 (3 islands) is one the adaptive arm’s own backbone screen rejected outright, culling all 16 diffused backbones, so it produced no sequence and no fold and its exclusion removes an adaptive failure. M0315 1ey3 (4 islands) is one the adaptive arm solved and the reference did not, so its exclusion costs the adaptive arm a discordant success. That loss is smaller than it appears. Every passing fold for that motif lies in the sixth branch of its lineage, so the solve arrives after roughly 800 of 960 fold predictions and at a redesign depth the reported configuration does not permit; including it would breach the matched 200-fold budget imposed on every other motif. Neither exclusion was chosen after seeing its effect on the test.

Two of the 36 matched motifs, M0129 1os7 and M0584 1ldm, reached the scheduler wall clock part-way through a step. They are scored as failures, so an incomplete adaptive run counts against the adaptive arm.

### GPU occupancy in packed small-molecule pipelines

**Figure S2:**
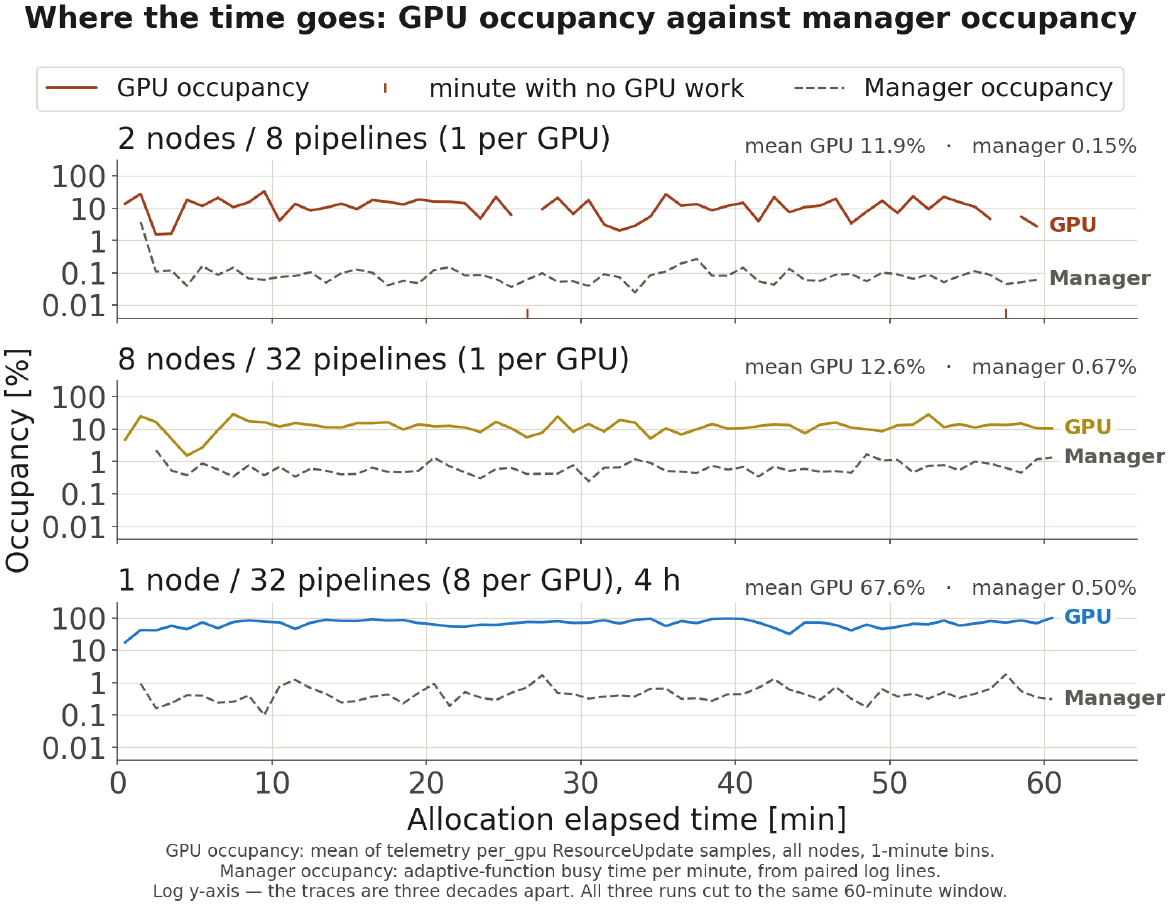
GPU occupancy in a 1-hour run measured in three pipeline loading conditions: 1) 8 pipelines, 1 per GPU; 2) 32 pipelines, 1 per GPU; and 3) 32 pipelines, 8 per GPU. Packing designs on the GPU led to significant performance gains over naive packing of 1 pipeline per GPU.

### Sequence-space pairing for the PDZ redesign campaign

**Figure S3:**
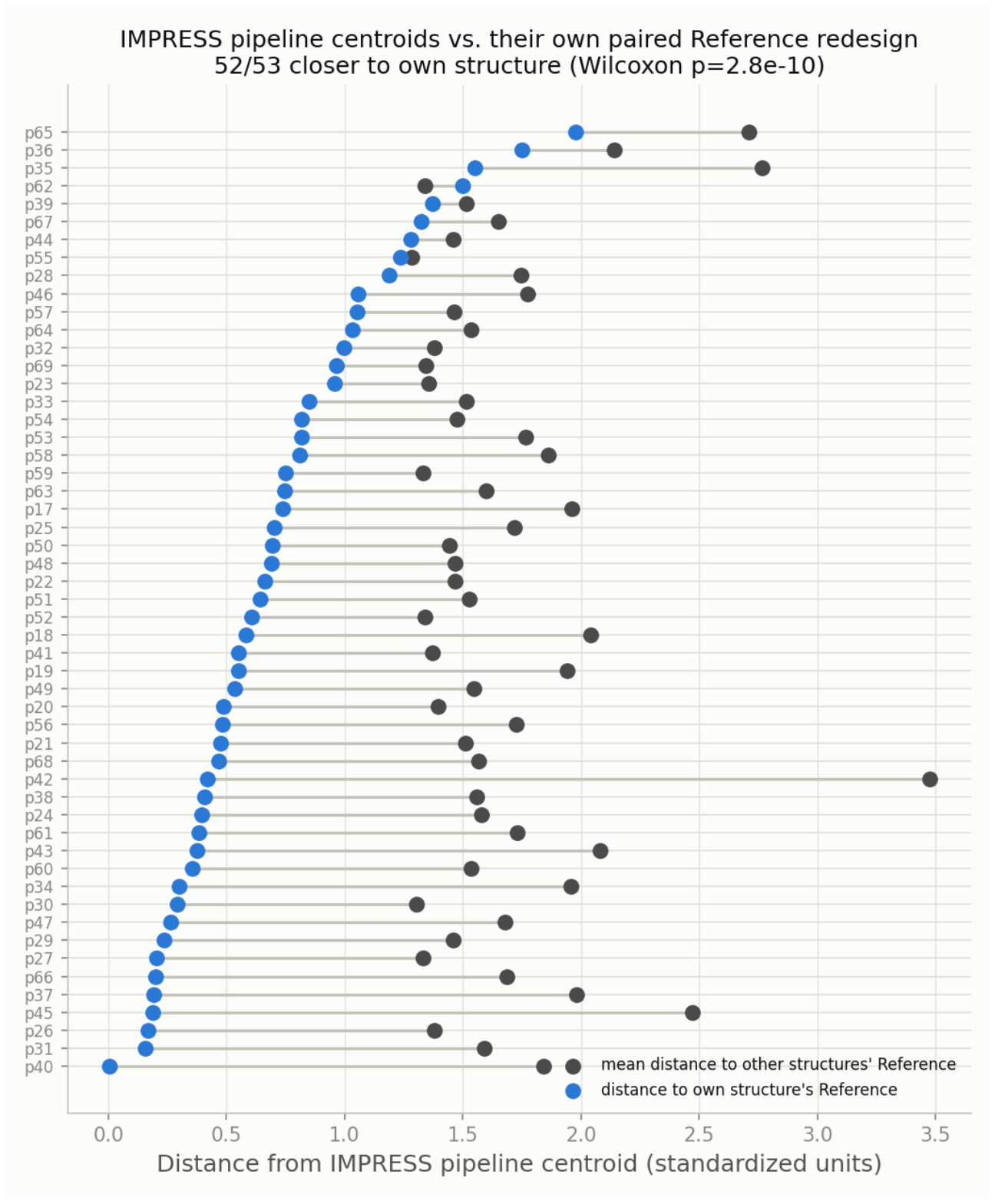
Adaptive PDZ redesigns stay near the input they started from. Each row is one of the 53 starting structures both arms redesigned; the quantity plotted is a distance in a standardized sequence space whose axes are per-residue amino-acid entropy and mean pairwise 2-mer cosine distance. For each structure the centroid of that pipeline’s adaptive sequences is compared with the nonadaptive redesign of the same structure (blue) and with the mean over the nonadaptive redesigns of all the other structures (grey). The adaptive centroid is closer to its own structure’s redesign for 52 of the 53 pipelines, a mean distance of 0.71 against 1.69 (Wilcoxon signed-rank *p* = 2.8 *×* 10*^−^*^10^). Two permutation tests on the same coordinates agree: a pipeline’s own sequences are tighter together than a randomly drawn group of the same size, 0.92 against a null mean of 1.84, and its centroid sits further from the pooled nonadaptive population than chance, 1.11 against 0.64 (*p* = 0.0002 each, 5,000 permutations). The input range, and a property of the pairing that bears on the effect size, are described above.

